# A cross-species spatial transcriptomic atlas of the human and non-human primate basal ganglia

**DOI:** 10.1101/2025.11.22.688128

**Authors:** Madeleine N. Hewitt, Meghan A. Turner, Nelson Johansen, Delissa A. McMillen, Shu Dan, Mike DeBerardine, Augustin Ruiz, Mike Huang, Jacob Quon, Yuanyuan Fu, Inkar Kapen, Stuard Barta, Naomi Martin, Nasmil Valera Cuevas, Paul Olsen, Josh Nagra, Jazmin Campos, Marshall M. VanNess, Shea Ransford, Zoe Juneau, Sam Hastings, Lindsey Ching, Michael Kunst, Soumyadeep Basu, Thomas Höllt, Chang Li, Boudewijn Lelieveldt, Faraz Yazdani, Qiangge Zhang, Kirsten Levandowski, Guoping Feng, Burke Q. Rosen, Matthew F. Glasser, Takuya Hayashi, Aaron D. Garcia, Omar Kana, Zoe M. Maltzer, Luke Campagnola, Tim Jarsky, Lauren Kruse, Winrich Freiwald, C. Dirk Keene, David C. Van Essen, Jeanelle Ariza, Jack Waters, Fenna M. Krienen, Trygve E. Bakken, Rebecca D. Hodge, Lydia Ng, Hongkui Zeng, Ed S. Lein, Jennie L. Close, Brian Long, Stephanie C. Seeman

**Affiliations:** Allen Institute for Brain Science, Seattle, WA; Princeton Neuroscience Institute, Princeton University, Princeton NJ; Leiden University Medical Center, Netherlands; Delft University of Technology, Netherlands; Laboratory of Neural Systems, The Rockefeller University, New York, NY; McGovern Institute for Brain Research, Department of Brain and Cognitive Sciences, Massachusetts Institute of Technology, Cambridge, MA; Stanley Center for Psychiatric Research, Broad Institute of MIT and Harvard, Cambridge, MA; Department of Neuroscience, Washington University in St. Louis, St. Louis, MO; Washington University School of Medicine, St. Louis, MO; Laboratory for Brain Connectomics Imaging, RIKEN Center for Biosystems Dynamics Research, Kobe, Japan; Department of Brain Connectomics, Kyoto University Graduate School of Medicine, Kyoto, Japan; University of Washington BioRepository and Integrated Neuropathology (BRaIN) lab, Harborview Medical Center, Seattle, WA

## Abstract

The basal ganglia are interconnected subcortical nuclei with complex topographical organization that orchestrate goal-directed behaviors and are implicated in neurodegenerative movement disorders. We generated a cellular-resolution, spatial transcriptomic atlas of the basal ganglia in human, rhesus macaque, and common marmoset, sampling over one million cells in each species. By integrating spatial data with a cross-species, consensus snRNA-seq cell type taxonomy, this atlas reveals conserved principles of molecular organization within and across structures. The cellular architecture is complex but highly stereotyped, with gene expression gradients superimposed onto discrete compartments. Extensive spatial sampling illuminates 3D gradients of molecular organization in the striatum and reveals cell type-specific core and shell compartments in the primate internal globus pallidus, which is conserved with mouse. This unified, cross-species spatial transcriptomic atlas will be a foundational resource for characterizing the molecular and functional organization of the basal ganglia and their roles in health and disease.

## Introduction

High-resolution transcriptional and anatomical maps of the mouse and human brain have laid the foundation for our current understanding of brain organization, connectivity, and cell type diversity^1–3^. Recently, single-cell transcriptomic technologies have refined these atlases, providing unprecedented detail of the cellular composition of the primate brain^4,5^. These resources have enabled the identification of conserved and derived species-specific features of cortical architecture and identified molecular markers for functionally distinct neuronal populations^6–10^. The simian to human lineage spans roughly 40 million years of evolution^11^, and while these existing studies have made significant progress describing cross-species cortical structure, the molecular conservation of subcortical areas remains under-characterized.

The basal ganglia are deep subcortical nuclei critical for motor control, decision-making, and reward processing that have yet to be comprehensively profiled and compared across primates. Canonical regions of the basal ganglia circuit include the striatum, globus pallidus, subthalamic nucleus, and substantia nigra^12^. Although the basal ganglia have been extensively studied in rodents^13–18^, including the generation of detailed anatomical and functional maps, key questions remain about how these structures are organized at the cellular and molecular level in primates, whose basal ganglia exhibit evolutionary divergence^15,19–23^. This knowledge gap is particularly significant given that non-human primates have advantages over rodents in studying various neuropsychiatric and neurodegenerative disorders that involve the basal ganglia (e.g., Parkinson’s disease, Huntington’s disease, and addiction^24,25^). Recent single-cell transcriptomic studies have begun to provide a detailed accounting of the cell types present in individual primate species in a subset of basal ganglia nuclei^26,27^. However, the tissue dissociation required for single-cell experiments limits their ability to link these molecular identities to the extensive, well-characterized topographic organization of the core basal ganglia nuclei. Providing spatial context to transcriptomic cell types in the primate, as is now readily possible via spatial transcriptomic techniques, is a critical missing link for understanding the functional organization of the basal ganglia.

Alongside other efforts that will be published concurrently^28^, we have generated a high-resolution, comprehensive, and multimodal basal ganglia cell type atlas across three primate species: marmoset, macaque, and human. The integrated atlas consists of three major data types that collectively classify transcriptomic cell types^29^, map their tissue distributions (this study), and characterize their properties^30^ within a harmonized ontological schema^31^. Together, these modalities provide a multifaceted view of primate basal ganglia cell types. In this study, spatial transcriptomics data were used to characterize cell type localization, explore gene expression gradients, and perform interspecies comparisons of spatial organization. Overall, we find highly conserved organization of basal ganglia nuclei among the three primate species that, in many cases, also extends to mouse. Cell types of each nucleus are organized into discrete domains, such as striatal striosomes or the core and shell of the internal globus pallidus. At the same time, gene expression gradients can span the full axis of the structure as exemplified in complex gene expression patterns along the internal capsule of the striatum. By integrating our data into online visualization platforms and analysis tools, we provide an accessible resource that can serve as a common framework for future basal ganglia research.

## Results

### A cross-primate spatial atlas of the basal ganglia

We present a cross-species spatial transcriptomic atlas of the basal ganglia, encompassing more than eight million cells across nearly 100 coronal sections (Fig 1A-C). We sampled striatum (STR), globus pallidus external (GPe) and internal (GPi) segments, substantia nigra (SN), and subthalamic nucleus (STH) in coronal sections that spanned the full rostral-to-caudal extent in each species (Supp Fig S1). Our experimental design accounted for the wide range of spatial scales across the three species with targeted sampling strategies. Marmoset sections were spaced approximately 200 µm apart while macaque and human sections were spaced approximately 1 mm apart (Fig 1A-C). We utilized the Vizgen MERSCOPE platform for human and macaque and the 10X Xenium platform for marmoset, each with gene panels comprising up to 300 genes. Gene selection was optimized for each species with at least 100 genes in common between each pair and 72 genes shared across all three species (Supp Table S1).

**Figure 1.**
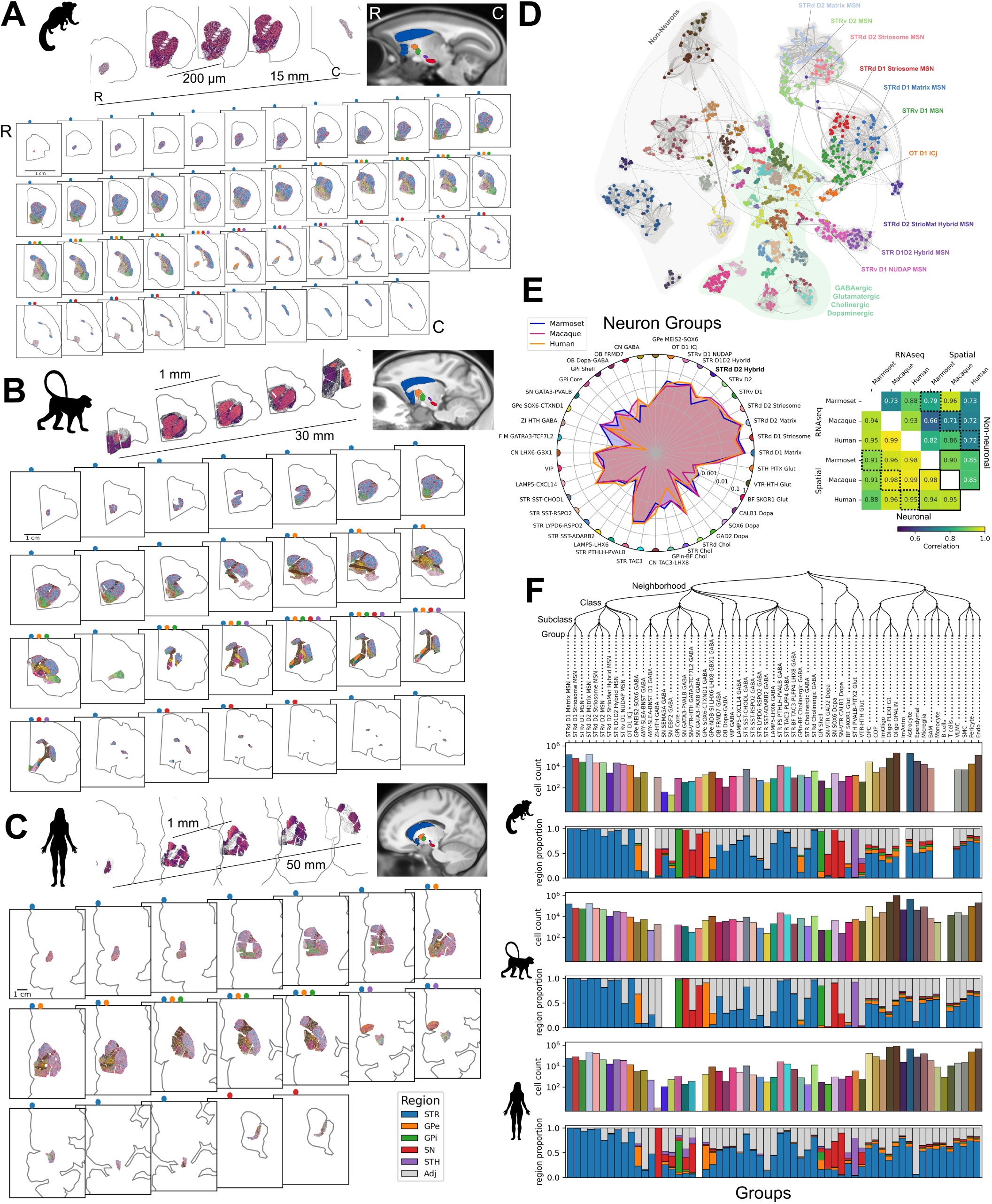
Cross species basal ganglia spatial transcriptomics atlas: **A-C.** Spatial transcriptomic sampling of the basal ganglia in three primates, common marmoset (A), rhesus macaque (B), and human (C). The top row depicts *DRD1* expression in spatial transcriptomics sections spanning the rostral (R) and caudal (C) ends of the sampling range and 3 middle sections depicting the sampling interval. To the left is a template sagittal view with BG CCF regions overlaid for reference. Below, all sections collected ordered from rostral to caudal with an outline of the full hemisphere. Spatial transcriptomic cells are colored by Group following colors in D and the color bars in F. Colored dots above each section indicate the BG regions sampled. **D.** UMAP of cross-species consensus taxonomy. Each dot is a cluster colored by Group. **E.** Comparison of Group proportions in spatial transcriptomics across species. The radar plot (left) includes all species, and each point is the number of cells in that Group as a fraction of all neuronal Groups plotted on a log scale. Heatmap (right) shows Pearson correlation of these populations across species and modality for neuronal (lower left half) and non-neuronal (upper right half) cells. Solid lines denote cross-species, within-spatial correlation; dotted lines denote within-species, cross-modality correlation **F.** BG transcriptomic hierarchy terminating at the species-conserved Group level (top). Each pair of plots below show cell count and region proportions for each Group in each species. Full Group names are used in F and short Group names in E for visibility (see SuppTable S2).

Several post-processing steps (Supp Fig S2) ensured comparable spatial transcriptomic datasets across species and platforms. For the MERSCOPE platform, cells were segmented using a custom Cellpose model^32^ (*Methods*); for the Xenium platform, cells were segmented using the cell segmentation staining kit and associated on-instrument algorithm (Supp Fig S2A). Following cell segmentation, cells with few transcripts or genes were removed (Supp Fig S2B, *Methods*) resulting in 1.3 million cells (96%) in marmoset, 3.1 million (97%) in macaque, and 4.0 million (76%) in human (Supp Fig S2D). We used MapMyCells (RRID:SCR_024672), a correlation-based mapping method, to assign each spatial transcriptomic cell a cell type label (Supp Fig S2C) from the Basal Ganglia Cross-Species Consensus Taxonomy^29^ (Fig 1D, Supp Table S2).

The consensus taxonomy comprises the neuronal and non-neuronal cell types of the basal ganglia and is hierarchically organized at five levels of increasing granularity: Neighborhood, Class, Subclass, Group, and species-specific clusters (Fig 1F, Supp Table S2). As described in Johansen et al., 2025, the Group level of the taxonomy (Fig 1D, dot colors) is the finest level of cell type resolution conserved across species. In this paper, we focus on the Group level to facilitate comparison of spatial distributions of cell types across species. After assigning consensus labels to spatial cells, we observe the full diversity of basal ganglia Groups in all three primate species, allowing us to perform parallel analysis of the spatial organization of these cell types (Fig 1F, multi-color bars; Supp Table S3-5). First, we compared the relative proportions of each Group across species (Fig 1E). Overall, Group proportions for both neuronal and non-neuronal Groups are highly correlated across species (Fig 1E right, solid outlines). We validate the snRNA-seq-based Group classification by comparing Group proportions between the two modalities (Supp Fig S3A). Neuronal Groups correspond well between spatial transcriptomics and snRNA-seq, whereas the proportions of non-neuronal Groups are less consistent (Fig 1E right, dotted outlines), likely because sorting of snRNA-seq cells depletes the proportions of non-neuronal cells^29^. Though there is general consistency across species of cell abundance among Groups, we find species differences in individual Groups. The newly described *STRd D2 StrioMat Hybrid* Group (Fig 1E, bolded) is a rare type that is uniformly distributed in the dorsal striatum and is three times as abundant in macaque and human compared to marmoset. In both the spatial (Fig 1E) and snRNA-seq data (Supp Fig S3A), this type represents approximately ∼1.5% of all neuronal cells in macaque and human versus 0.5% in marmoset (Supp Table S6).

Many neuronal Groups show regional localization, such as medium spiny neurons (MSNs) predominantly found in the striatum, in contrast to non-neuronal Groups with more ubiquitous coverage (Fig 1F, Supp Table S7). Inhibitory interneuron Groups of the MGE and CGE lineage show varied regional distributions (Supp Fig S4-S6). Our data confirm that *TAC3-*expressing interneurons^27,33^ in the *STR TAC3-PLPP4 GABA* Group are localized almost exclusively to the striatum in all three species (proportion in STR: marmoset, 0.9; macaque, 1.0; human, 0.9).

Additionally, we identify a second *TAC3* and *LHX8* expressing Group, *STR-BF TAC3-PLPP4-LHX8 GABA*, in both the striatum and surrounding areas (proportion in STR: marmoset, 0.6; macaque, 0.7; human, 0.7). We also find regional specialization in the cholinergic Groups.

*STRd Cholinergic GABA* cell localization is biased to the dorsal striatum, while *STR Cholinergic GABA* cells localize to the ventral striatum and basal forebrain (Supp Fig S4-S6). *GPin-BF Cholinergic* cells are found specifically in the lamina surrounding GPe and GPi as well as in the basal forebrain. Consistent sampling through the rostral-caudal (R-C) axis of the basal ganglia allows for a deeper inspection of cell type localization through these structures (Supp Fig S3B). For instance, in macaque and marmoset, peak *OT D1 ICj* cell density is in the most rostral section and is highly localized, whereas human *OT D1 ICj* cells localize more caudally and are more diffuse, consistent with previous descriptions of the olfactory tubercle (OT)^19,34–36^. D1 and D2 MSNs of the dorsal striatum are broadly distributed through the R-C axis but with a rostral bias in all three species.

### Cross-species Basal Ganglia Atlas Resources

We have incorporated our spatial transcriptomics atlas into two interactive platforms to enable exploration of basal ganglia cell types and gene expression across species. First, these data have been integrated into the Allen Institute’s ABC Atlas (RRID:SCR_024440), a web-based platform for visualization of cellular data^10^. The ABC Atlas allows users to view spatial transcriptomic data from all three primate species alongside an integrated cross-species embedding of the snRNA-seq consensus taxonomy, with interactive selection and filtering. In addition to cell types, users can explore gene expression and continuous variation across the basal ganglia in all three species simultaneously. Second, the data from this paper and others have been incorporated into the Cytosplore Viewer desktop application^37^ and its Gradient Surfer plugin (Supp Fig S7), which offers interactive analyses to compare spatial gene expression gradients across two datasets with partially overlapping gene panels. It is designed to interactively probe for, extract, and align gradient-based gene expression features to identify conserved or divergent molecular patterns related to tissue structure and biological function.

Additionally, the data used in this paper are available for download alongside tutorials for programmatic access (https://brain-map.org/consortia/hmba/hmba-release-basal-ganglia).

### Discrete organization of striatal neurons

The striatum (STR)—consisting of the caudate, putamen, nucleus accumbens, and olfactory tubercle—is the primary input nucleus of the basal ganglia. The primate striatum is discretely organized in two intertwined ways: distinct transcriptomic profiles (Fig 2A) and neurochemically defined compartments (Fig 2B). The four neurochemically defined compartments are striosomes and the surrounding matrix, Islands of Calleja (ICjs), and <u>n</u>eurochemically <u>u</u>nique <u>d</u>omains found in the nucleus <u>a</u>ccumbens and <u>p</u>utamen (NUDAPs). While these compartments were originally defined by neurochemical or histochemical staining^38,39^, modern molecular techniques have identified unique transcriptomic cell types associated with each compartment^17,26^. The consensus taxonomy identifies transcriptomically defined Groups associated with each of these compartments (Fig 2A): *STRd D1 Striosome MSN* and *STRd D2 Striosome MSN* in striosomes; *STRd D1 Matrix MSN* and *STRd D2 Matrix MSN* in matrix; *STRv D1 NUDAP MSN* in NUDAPs, and *OT D1 ICj* in ICjs. We used the Group labels on our spatial cells to generate polygon masks of the striosome, NUDAP, and ICj compartments (Fig 2B, *Methods*). Approximately 12% of the total striatal area sampled in our coronal sections belongs to one of these three compartments, with the remaining area comprises the matrix compartment. The striosomes dominate the compartment fraction, covering 10%-11% of total striatal area per species (Fig 2C, right). In the marmoset, the NUDAPs and ICjs represent a larger proportion of the striatum than in the other two species, which may reflect the expansion of the ventral striatum in marmoset.

**Figure 2.**
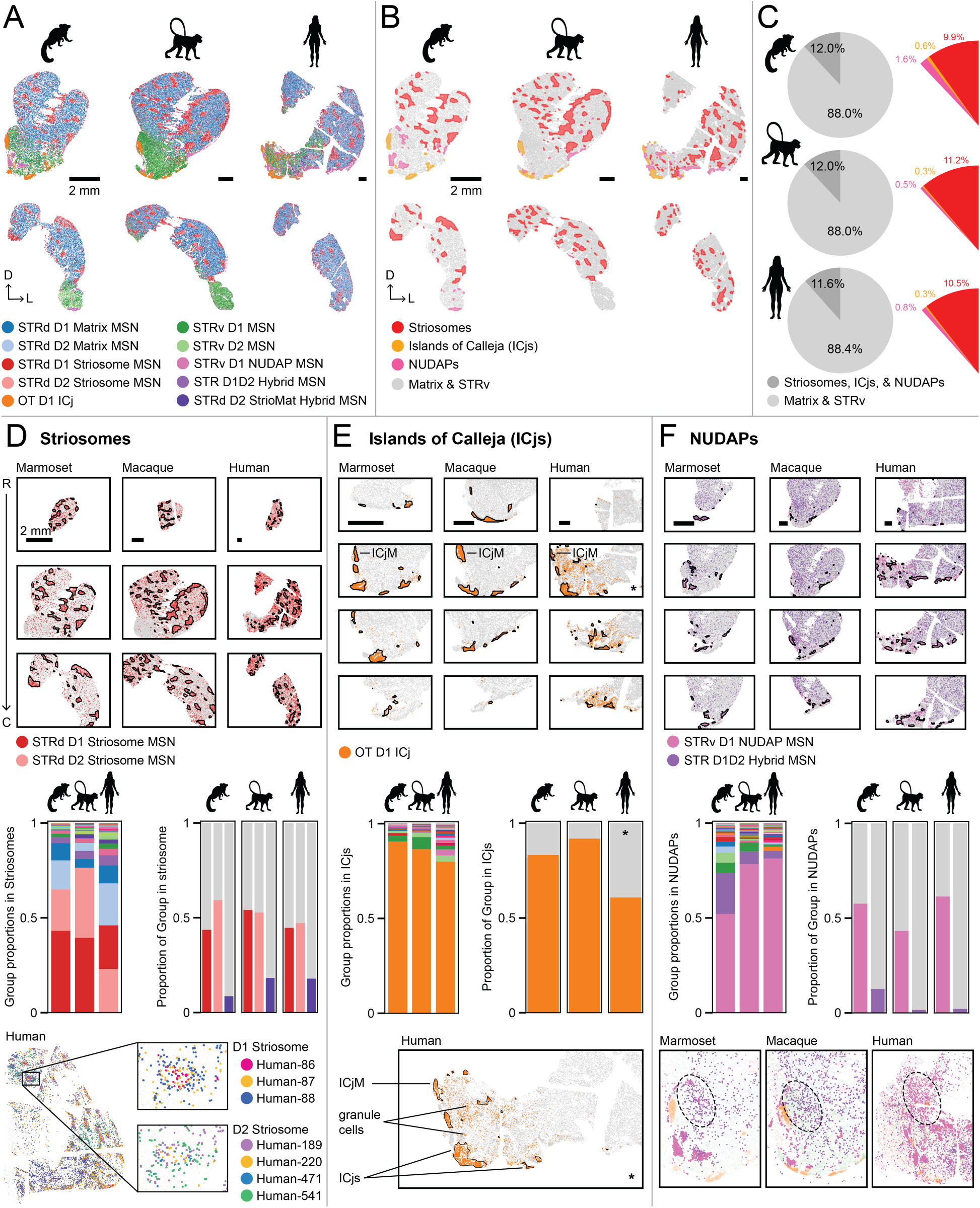
Discrete cell types and compartments of the striatum. **A.** The ten predominant neuronal Groups in the striatum, shown at one rostral and one caudal z-plane in each species. **B.** Polygon masks of three compartments: striosomes (red), ICjs (orange), and NUDAPs (pink) in the same sections as in A; all scalebars shown are 2 mm. **C.** Left, fraction of striatal neurons that are in a striosome, ICj, or NUDAP compartment (dark grey) vs the matrix or non-compartment STRv (light grey). Right, fraction of striatal neurons that belong to each of the striosome, ICj, or NUDAP compartments. **D.** Striosomes: (Top) Outlines of striosome compartment masks overlaid on *STRd D1 Striosome MSN* and *STRd D2 Striosome MSN* cells. (Middle, left) Proportions of striatal Groups found inside striosome masks. (Middle, right) Proportion of cells from each of three Groups (*STRd D1 Striosome MSN, STRd D2 Striosome MSN, STR StrioMat Hybrid*) that are located inside (red) vs outside (grey) striosome polygon masks. (Bottom) Representative striosome, in human, exhibiting an annulus and core spatial organization of both *STRd D1 & D2 Striosome MSN* clusters **E.** ICjs: (Top) Outlines of ICj compartment masks overlaid on *OT D1 ICj* cells. (Middle, left) Proportions of striatal and interneuron Groups found inside ICj masks. (Middle, right) Proportion of cells from the *OT D1 ICj* Group that are located inside (orange) vs outside (grey) striosome polygon masks. (Bottom) Single z-plane in human showing the three types of localizations of granule cells: in the ICjM, in canonical ICjs, and scattered throughout the STRv. **F.** NUDAPs: (Top) Outlines of NUDAP compartment masks overlaid on *STRv D1 NUDAP MSN* and *STR D1D2 Hybrid MSN* cells. (Middle, left) Proportions of striatal Groups found inside NUDAP masks. (Middle, right) Proportion of cells from each of two Groups (*STRv D1 NUDAP MSN* and *STR D1D2 Hybrid MSN*) that are located inside (pink, purple) vs outside (grey) NUDAP polygon masks. (Bottom) Example sections from each species showing the diffuse mixture of *STRv D1 NUDAP MSN* and *STR D1D2 Hybrid MSN* Groups localized to the NACsmd.

The spatial atlas allows us to disambiguate each Group from its namesake compartment and quantify the overlap between the two. For example, we identify cells mapping to striosome Groups outside of the boundaries of the striosome compartments in all three species (Fig 2D, top). Using the striosome compartment masks, we find that approximately half of all *STRd D1 Striosome MSN* and *STRd D2 Striosome MSN* cells in each species lie outside of the striosome compartments (Fig 2D, middle). These may be a primate homolog of the “exo-patch” cells reported in mouse, which are scattered throughout the matrix but molecularly resemble MSNs found in striosome compartments^40,41^. Notably, we find a significantly higher proportion of putative exo-patch cells (∼50%) than the current estimates in mouse (12%)^42^. Additionally, our results reveal spatial organization among the transcriptomic cell types within striosome compartments. In human striosomes, clusters from both the *STRd D1 Striosome MSN* and *STRd D2 Striosome MSN* Groups form overlapping radial layers; specific clusters are biased to an inner “core” zone while others are biased to an outer “annulus” zone (Fig 2D, bottom), revealing a cellular analog to histochemical features observed in human striosomes^43^.

In all primate species, we find that granule cells (*OT D1 ICj* Group) localize into three previously described domains^44^ (Fig 2E): (1) the insula magna of Calleja (ICjM), a stereotyped island at the medial interface between the nucleus accumbens and basal forebrain; (2) canonical islands (ICjs), which are distributed across the olfactory tubercle; and (3) smaller clusters and individual cells scattered throughout the ventral striatum and basal forebrain structures. This spatial atlas confirms that the proportion of *OT D1 ICj* cells that localize to the third domain in human is significantly higher than in marmoset or macaque^34^.

We also identified an MSN Group associated with NUDAP compartments, sometimes referred to as “interface islands”^26,45^, in all three species (Fig 2F, top). We find a range of sizes, shapes, and cellular densities across the NUDAP compartments. Notably, around half of *STRv D1 NUDAP MSN* cells reside outside the boundaries of the NUDAP compartments (Fig 2F, middle). For example, there is a subpopulation of *STRv D1 NUDAP MSN* cells that do not fall into a NUDAP compartment but are enriched in the mediodorsal subdivision of the shell of the nucleus accumbens (NACsmd) (Fig 2F, bottom). We also find another cell type that localizes to the NUDAP compartments, the *STR D1D2 Hybrid MSN* Group, which illustrates that there is not always a one-to-one relationship between neurochemical compartments and molecular cell types.

### Continuous variation of gene expression in the dorsal striatum

Superimposed on the striatum’s discrete substructure is a graded functional organization composed of overlapping motor, associative, and limbic areas in a dorsolateral-to-ventromedial (DL-VM) fashion^12,15,19,20,46,47^. We observed smoothly varying gene expression along this same axis within the MSNs of the dorsal striatum (STRd). This variation is primarily observed as a gradient of gene expression that increases or decreases along the direction of the internal capsule in coronal sections through the caudate and putamen (Fig 3, Supp Fig S9). This gradient has been documented in both rodents and primates and is typically illustrated by increasing *CNR1* and decreasing *CRYM* expression along the ventromedial-to-dorsolateral (VM-DL) axis of the striatum^14,17,18,26,40^. Although measured expression levels are different in each species, we observed that the *CNR1* gradient is conserved across species and present in multiple coronal planes (Supp Fig S9). To identify other genes that covary with *CNR1,* we performed non-spatial Principal Component Analysis (PCA) on the gene expression of all cells in the *STRd MSN* Groups in each species. We selected the PC with the highest correlation to the internal capsule axis, which we refer to as the Principal Gradient Component of the dorsal striatum, “PGCd” (PC3, PC2, and PC4 for marmoset, macaque, and human). The PGCd represents an important organizing feature in the striatum. The PGCd gradient direction in the coronal plane is conserved across D1 and D2 striosome and matrix MSNs (Fig 3A). While the average PGCd changes across the rostral-caudal axis, this pattern is highly correlated across the four MSN Groups (Fig 3B). This conserved gradient includes additional genes beyond the canonical *CNR1*, such as *TESPA1* and *GDA*, that reflect variation along the PGCd (Fig 3C, Supp Fig 9C). With the Gradient Surfer tool in the Cytosplore Viewer desktop application (Supp Fig S7), described above, researchers can explore the continuous variation of gene expression in STRd.

**Figure 3.**
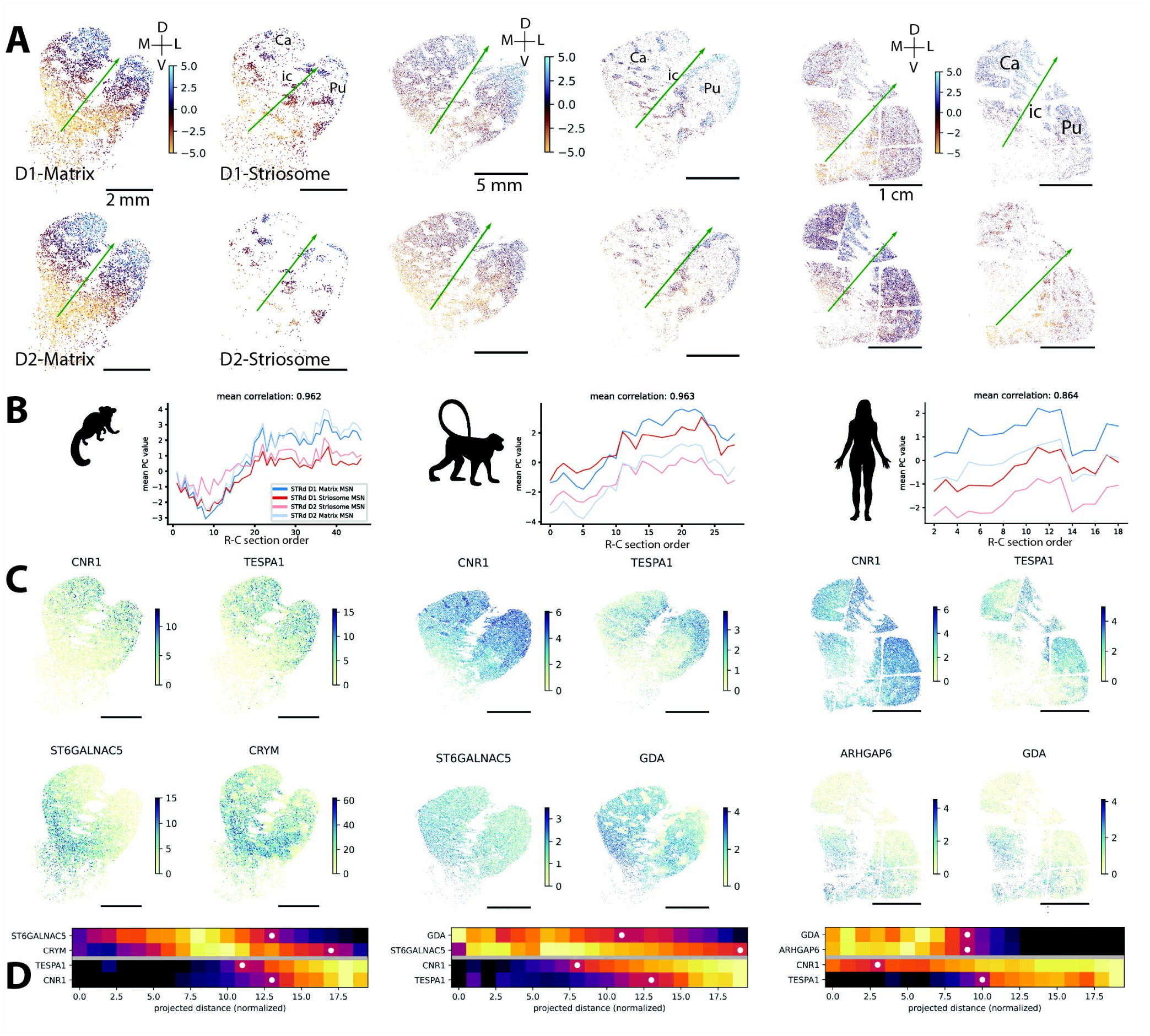
Gradient organization of the dorsal striatum: **A.** Principal component analysis of STRd MSN Groups showing the dorsal principal gradient component (PGCd) in marmoset, macaque and human. The arrow indicates the PGCd direction calculated from the cells of each Group. (Abbreviations: Ca: Caudate, ic: internal capsule, Pu: putamen). **B.** Average PGCd values for each group across rostral-caudal (R-C) order are highly correlated with each other. **C.** Gene expression in each species showing examples of genes that increase (*CNR1*) and genes that decrease (*GDA*) in expression towards the dorsal-lateral direction. **D.** Heatmaps showing average gene expression vs the projected distance along the internal capsule axis (see Supp Fig S9B) for genes in the PGCd. White dots indicate the half-maximum position along this coordinate. Even though *CNR1* and *TESPA1* are correlated with each other and change in the same direction along the gradient in each species, their spatial distributions are distinct, characterized here by the different half-maximum positions. Further examples of smoothly varying gene expression in the PGCd are shown in Supplemental Figure S9C.

We analyzed the expression of the genes contributing to the PGCd pattern and found that while they do vary along this gradient, their spatial patterns include more independent variation than simply increasing or decreasing in lockstep with *CRYM* or *CNR1* (Supp Fig S9C). To quantify this and characterize the pattern of each gene independently, we projected gene expression along the gradient axis and identified the points where gene expression was half of the maximum value. These half-maximum points vary across genes (Fig 3D), showing that the relationship between gene expression and VM-DL position differs across genes. Previous analyses of these gradients in mouse have suggested either a single axis of variation defined by the *CNR1*/*CRYM* ratio^17^ or two opposing gene programs^18^. Our data show that the PGCd in these three species is a combination of diverse gene gradients that do not all share an identical spatial pattern. This high degree of spatial diversity in MSN gene expression may reflect complex functional connectivity in striatal circuits.

### Continuous variation of gene expression in the ventral striatum

The ventral striatum (STRv) is a histochemically heterogeneous region that includes the nucleus accumbens (NAC), which can be further divided into a core (NACc) and shell (NACs). *CALB1,* or its protein product calbindin, is often cited as a definitive marker for NACc and NACs, but this is not consistent across species or rostral-caudal positions within a species^48^ (Supp Fig S10).

Previous studies have also noted a lack of histochemical or molecular markers that distinguish NACc and, more broadly, STRv from STRd^17,23,26,49^. This ambiguity agrees with the observation of Ding et al. 2025^31^ [concurrently published] that different marker genes (e.g., *CALB1, WFS1*) suggest different boundaries between dorsal and ventral striatum. We therefore sought to identify gene expression patterns that define or characterize the ventral striatum while also considering its surrounding anatomical context.

The consensus taxonomy distinguishes between four STRd MSNs Groups, discussed above, and two STRv MSN Groups that show similar localization along the R-C axis. In the rostral striatum, STRv MSNs are found primarily in the NAC and OT (Fig 4A), which are also characterized by pockets of higher ratios of D1 to D2 MSNs (Fig 4B). To identify genes associated with the transition from dorsal to ventral striatum, we expanded our PCA decomposition to include *STRv D1 MSN, STRv D2 MSN*, and *AMY-SLEA-BNST GABA* Groups in addition to the STRd Matrix and Striosome Groups. We included *AMY-SLEA-BNST GABA* because many genes expressed in STRv extend smoothly into the bed nucleus of the stria terminalis (BNST) and central nucleus of the amygdala (CEN). Similar to the PGCd, for each species, we created a line approximating the internal capsule axis, extending this line into the ventral striatum. We then selected the PC that had the highest correlation to this axis, which we refer to as the PGCv. Overall, the PGCv is oriented similarly to PGCd, progressing smoothly in a DL-to-VM direction parallel to the internal capsule. However, within the STRv, the PCGv changes orientation and progresses medially towards the mediodorsal NAC shell (NACsmd).

**Figure 4.**
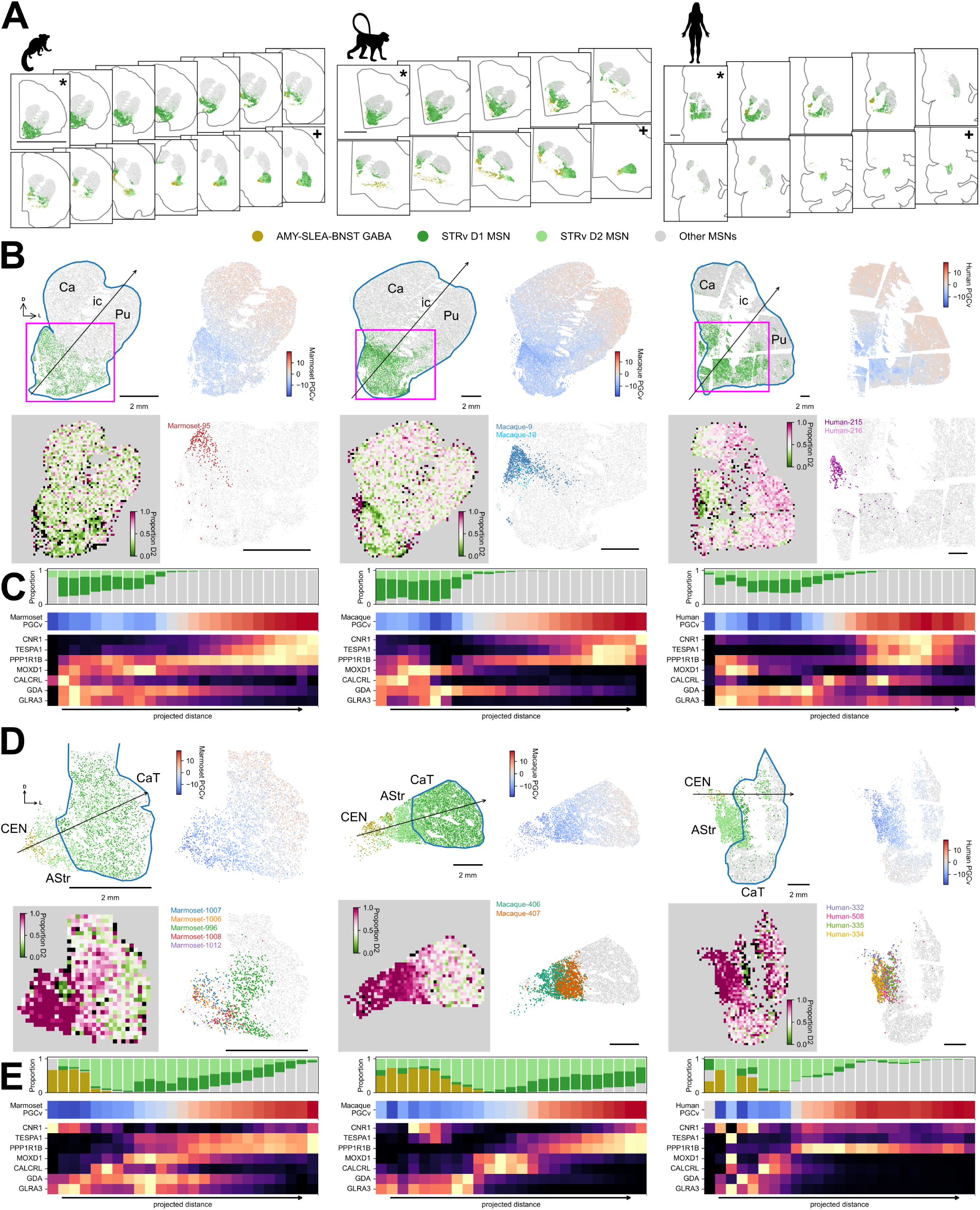
Continuous variation in the ventral striatum: **A.** Selected sections showing the rostral-to-caudal extent of *STRv D1* and *D2 MSNs* in marmoset (left), macaque (middle) and human (right). Asterisk (*) indicates the section analyzed in B and C. Plus sign (+) indicates the section analyzed in D and E. Abbreviations: Ca: Caudate, ic: internal capsule, Pu: putamen. **B.** For each species, select MSN Groups (top left), the PGCv (top right), spatial histograms of the proportion of D2 MSNs (bottom left), and selected NACsmd clusters (bottom right) are shown for a representative rostral section. Cluster plots are cropped to the box (magenta) in the top left plot for each species. **C.** For each species, cell type proportions, PGCv scores, and select genes are plotted for cells projected along the line in B and binned. Rows in the gene expression histogram are min-max normalized. **D.** For each species, select MSN Groups (top left), the PGCv (top right), spatial histograms of the proportion of D2 MSNs (bottom left), and selected AStr clusters (bottom right) are shown for a representative caudal section. Abbreviations: CaT: caudate tail, AStr: amygdalostriatal transition area, CEN: central nucleus of the amygdala. **E.** For each species, cell type proportions, PGCv scores, and select genes are plotted for cells projected along the line in D and binned. Rows in the gene expression histogram are min-max normalized.

The NACsmd is a triangular region of STRv below the ventricle that has previously been shown to have unique connectivity and molecular properties^50–53^. In this spatial atlas, each species has one or two *STRv D1 MSN* clusters that localize specifically to the NACsmd (*Marmoset-95*, *Macaque-9* and *10*, *Human-215* and *216*) and have lower *PPP1R1B* expression than other *STRv D1 MSNs* (Supp Fig S11).

To see how genes vary along the DL-VM axis in rostral striatum, we projected the cells onto the internal capsule axis line. To identify shared genes across species with graded or stepwise changes along this axis, we selected genes that contributed highly to the PGCv; some genes were also selected by manual inspection. The sign of the PGCv changes at the position along this axis where STRd types decrease in abundance and *STRv D1* and *D2 MSNs* increase, supporting its association with ventral cell type identity (Fig 4C). Some genes that show smooth gradients with lower expression in the ventral part of the dorsal striatum, such as *CNR1* and *TESPA1*, are also expressed in subregions of STRv (Supp Fig S11). Other genes are expressed in specific discrete locations along the PGCv, some of which are conserved across species (e.g., *MOXD1*) while others show species-dependent localization (e.g., *CALCRL*).

Notably, we do not observe a set of genes, in any of the three species, that consistently distinguishes STRv from STRd or NACs from NACc. There is not a simple molecular signature that defines ventral from dorsal striatum in our gene panel, but rather a suite of genes with varied expression patterns whose combinations mark different territories of the ventral striatum.

Regions of the caudal striatum, including caudate tail (CaT) and caudoventral putamen (PuCv), have limbic connectivity and similar molecular properties to rostral STRv regions such as NAC^19,54,55^. Rather than being constrained solely to NAC and OT, we find *STRv D1* and *D2 MSNs* at a similar ventromedial position across the longitudinal axis of the Ca (CaH, CaB, and CaT), in a narrow medial strip of caudal putamen, and in PuCv (Fig 4A), suggesting molecular cell types may align with limbic domains in caudal striatum. Across all species, we observe a region between the striatum and central nucleus of the amygdala that contains MSNs. This region is characterized by weaker *PPP1R1B* expression (Supp Fig S11) and a high proportion of D2 MSNs (compared to D1) (Fig 4D). We identify this region as the amygdalostriatal transition area (AStr), which is D2 MSN-enriched in mouse^56^ and shares histochemical features with NAC in primates^19^. Compared to the rostral striatum, the PGCv exhibits similar spatial variation in the caudal striatum, progressing DL-to-VM from lateral CaT through AStr and CEN. Several *STRv D2 MSN* clusters are unique to AStr (*Marmoset-996, 1006, 1007, 1008*, and *1012; Macaque-406* and *407*; *Human-332, 334, 335*, and *508*) and express less *PPP1R1B* than neighboring striatal MSNs (Fig 4D, Supp Fig S11). Along an axis drawn from CEN to dorsolateral CaT/PuCv, genes that have graded or discrete expression patterns in rostral STRv often exhibit similar features in CaT and AStr (Fig 4E, Supp Fig S11). This suggests shared gene expression patterns in rostral and caudal ventral striatum, and more specifically highlights NACsmd and AStr as ventral striatum subregions with unique molecular features. The spatial context provided by this atlas highlights these unique features, particularly in the AStr which has rarely been profiled^56^.

### Molecular organization of the subthalamic nucleus

The subthalamic nucleus (STH) contains overlapping functional domains along its longitudinal, VM-DL axis: a motor region in the dorsolateral third, a limbic region at the ventromedial tip, and an associative region that forms a transition region between the two^57,58^. Recent molecular studies in the mouse and macaque show that the molecular organization of the STH parallels these topographical gradients of connectivity^59–62^. We find that marmoset and human also share these key principles of spatial organization in STH cell types and gene expression. The cells of the STH predominantly belong to a single, glutamatergic Group in our consensus taxonomy, *STH PVALB-PITX2 Glut* (Fig 5A), in line with the identity of STH as a primarily excitatory structure^63^. Within this Group, we identified four or five unique clusters per species (Supp Fig S12). In each species, two predominant clusters subdivide the STH along its longitudinal axis, with some overlap in the center: *Marmoset-1469* (VM-biased) and *Marmoset-1470* (DL-biased) (Fig 5B, left); *Macaque-474* and *Macaque-471* (Fig 5B, center); *Human-395* and *Human-549* (Fig 5B, right). In our macaque data, a third cluster, *Macaque-473*, forms a distinct cap at the medio-ventral end of the STH (Fig 5B, center).

**Figure 5.**
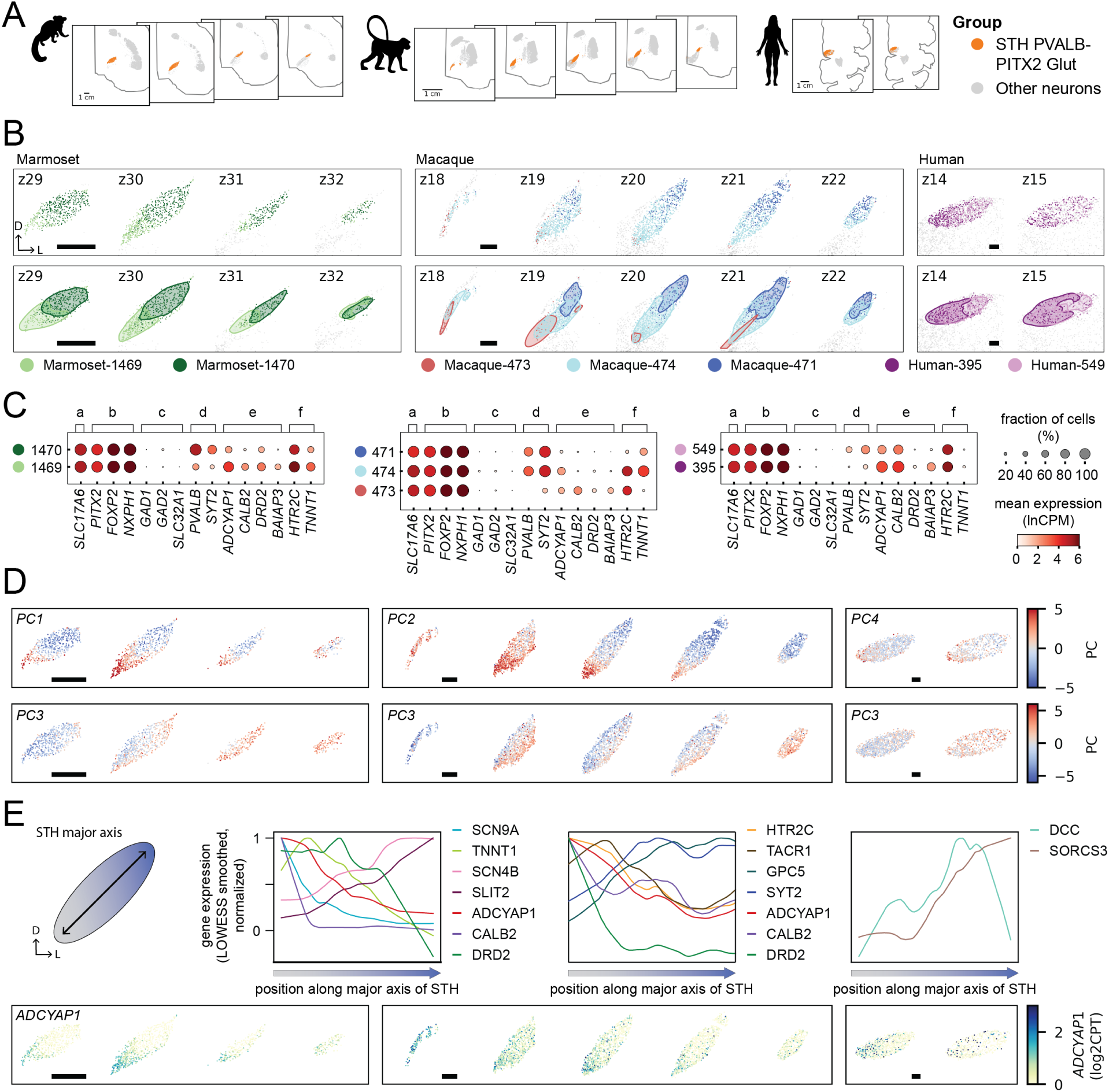
Spatial organization of the subthalamic nucleus. **A.** Montage of coronal sections containing STH, from rostral to caudal, in (left to right) marmoset, macaque, and human, with *STH PVALB-PITX2 Glut* Groups cells in orange and other cells colored grey. All scalebars are 1 mm. **B.** Dominant clusters in the STH (top) and the Kernel Density Estimate (KDE) contour containing 80% of the cells in each cluster (bottom) showing their biased spatial distribution along the VM-to-DL axis. **C.** Differential gene expression in the snRNA-seq data for clusters shown in B. **D.** PCA showing the two conserved axes of spatial variation in each species: longitudinal (top), and transverse plus rostral-caudal (bottom). **E.** Spatially variable gene expression in each species along the longitudinal, or major, axis of the STH. Each species has unique genes (top), but in all three species, *ADCYAP1* is high in the VM and low in the DL subthalamic nucleus (bottom).

Previous work in mouse and macaque have reported a conserved, glutamatergic *PVALB+* subpopulation restricted to the dorsolateral region of the STH^62,64–66^. The consensus taxonomy identifies glutamatergic *PVALB+* subpopulations (*Marmoset-1470*, *Macaque-471*, and *Human-549*), which are spatially biased to the dorsolateral end of the STH axis. The VM-biased clusters (*Marmoset-1469*, *Macaque-474*, and *Human-395*) also exhibit some *PVALB* expression, albeit in a smaller fraction of cells and with lower mean expression, consistent with overlap between these subpopulations.

To quantify the spatial axes of gene expression, we performed per-species, non-spatial PCA on the gene expression of all cells belonging to the *STH PVALB-PITX2 Glut* Group. We find two PCs corresponding to spatial axes that are conserved across all three primate species: (1) a longitudinal component that aligns with the ventromedial-to-dorsolateral axis highlighted by the cell type clusters (Fig 5D, top; Fig 5E), and (2) a transverse component that varies across the rostral-caudal axis and the minor (dorsomedial-to-ventrolateral) axis in sections collected near the midpoint of the rostral-caudal axis (Fig 5D, bottom). These axes of molecular variation are driven by partially overlapping sets of genes in the three species. All three species show graded expression of *ADCYAP1* along the longitudinal axis of STH (Fig 5E), with high expression at the ventromedial end decreasing to minimal expression at the dorsolateral end. *CALB2* exhibits a similar gradient in marmoset and macaque. We also find species-specific genes that share this high-VM/low-DL gradient pattern (Fig 5C,E; Supp Fig S13): *SCN9A*, *TNNT1*, *SCN4B* and *SLIT2* in marmoset; and *HTR2C*, *TACR1*, and *GPC5* in macaque.

We identified several differences across the three species. In macaque only, a third cluster, *Macaque-473*, forms a cap at the medio-ventral end of the STH (Fig 5B, center). Projection studies indicate that the medial tip has distinct properties from the rest of the medial half of the STH in macaque^57^. Furthermore, there is significantly more overlap between the two dominant clusters in human (Fig 5B, right) than NHPs. This is consistent with imaging studies that found more overlap between functional domains and more inter-individual variation in human STH^58,67^. Finally, while we find similar axes of transcriptomic diversity in primate as previously reported in mouse, the marker genes that drive these gradients are frequently unique to a single species. One notable example is *TAC1*, a pan-STH marker in the marmoset (Supp Fig S13), but a marker for just the neighboring parasubthalamic nucleus (pSTH) in mouse^68^. Despite these species differences, our spatial atlas demonstrates that the overall molecular organization of STH along its longitudinal axis is conserved across primate and mouse.

### Cell type organization of the Globus Pallidus

The globus pallidus (GP) is a central hub of the basal ganglia that integrates inhibitory and excitatory signals from the striatum and subthalamic nucleus to regulate motor output and motivational or limbic processing^69,70^. The GP includes the external segment (GPe) and internal segment (GPi), each with distinct molecular and functional identities. We find six Groups in the GP: two in the GPi, three in the GPe, and one Group of cholinergic cells in and around the GPe and GPi (Fig 6A).

**Figure 6.**
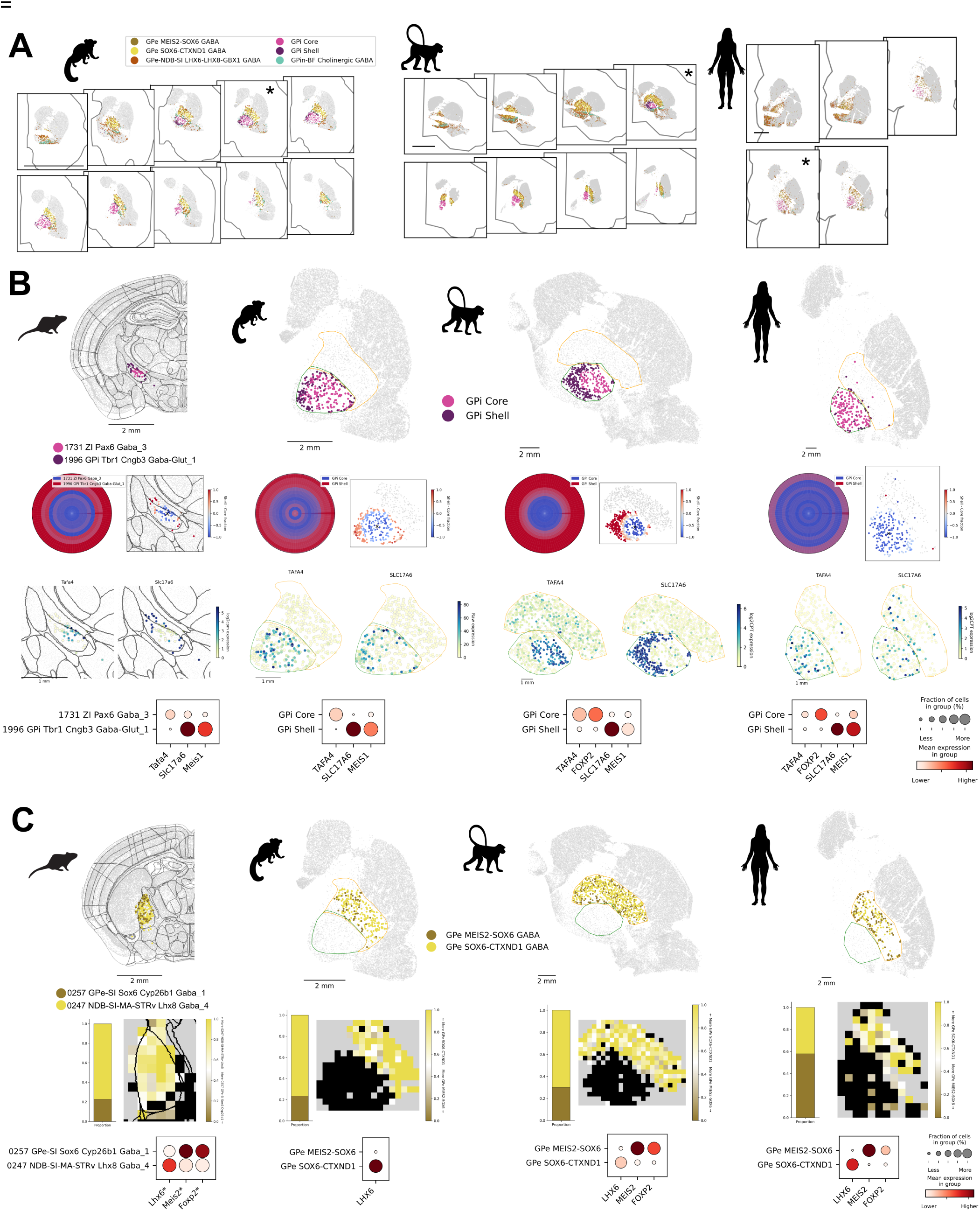
Cell type localization in the globus pallidus: **A.** Montage of coronal sections containing GP, from rostral to caudal, in each species with GP cells colored by Group and other cells colored grey. Sections denoted with * are highlighted in B and C. Scalebar represents 1cm. **B.** Top row shows a single slice from mouse, marmoset, macaque, and human with *GPi Core* and *Shell* Groups in primates and homologous mouse clusters. Middle row: Radial histogram (left) of GPi types with projection into spatial coordinates (right). Bin size (in microns) for mouse is 50, marmoset 100, macaque 200, and human 600. Center is denoted by an X. Bottom row: Gene expression of *TAFA4* and *SLC17A6*, marker genes for *GPi Core* and *Shell* respectively shown in spatial coordinates and as expression dot plots. **C.** Top row: Single slice with *GPe MEIS2-SOX6 GABA* and *GPe SOX6-CTXND1 GABA* Groups in primates and homologous supertypes in mouse. middle: Relative proportions of these two GPe types (left) and spatial histograms (right). Yellow/brown areas are the fraction of each Group in each bin on a diverging color scale while black bins do not contain either of these Groups. Bin size (in microns) for mouse is 150, marmoset 200, macaque 350, and human 800. Bottom row: Expression dot plots of marker genes. *Note that mouse expression data are imputed values. *FOXP2* and *MEIS2* were not in the marmoset gene panel.

The mouse homolog to GPi, the entopeduncular nucleus, was recently shown to have a core-shell division in its circuitry, which is mirrored in molecular cell types^71,72^. We find that the two GPi Groups in primate follow the same spatial localization as in mouse, with the *GPi Core* Group restricted to a central core surrounded by a band of *GPi Shell* Group cells. Through integration of RNA-seq-based taxonomies, we identified potentially homologous cell types in mouse^10^ for both subpopulations: mouse cluster *1731 ZI Pax6 Gaba_3* as a homolog to the primate *GPi Core* Group and mouse cluster *1996 GPi Tbr1 Cngb3 Gaba-Glut_1* as a homolog to the primate *GPi Shell* (Fig 6B). Radial histograms of *GPi Core* to *GPi Shell* proportions highlight the boundary between these subdivisions in mouse, marmoset, and macaque. In human, the *GPi Shell* represents a narrower band of cells, perhaps reflecting limited R-C sampling of this relatively small substructure. Gene expression from both the spatial transcriptomics data and the accompanying snRNA-seq data demonstrate high conservation across species (Fig 6B, Supp Fig S14A). For example, *TAFA4* is expressed in the *GPi Core* Group in all three primate species and mouse; *SLC17A6* and *MEIS1* are similarly conserved in the *GPi Shell* Group (Fig 6B). Expression of *FOXP2* is a hallmark of the *GPi Core* groups in all species (Supp Fig S14A). Here, for the first time, we show that distinct transcriptomic types in the primate GPi are organized into a core-shell structure and that these types have molecular and spatial homologs in mouse, suggesting a common organization of cell types may underly conserved differences in cell type projection targets across these species.

The GPe sends topographical projections to the STH in addition to GPi, SN, thalamus, and the pedunculopontine nucleus^24,70,73,74^. We find two GPe-specific Groups in primate, *GPe SOX6-CTXND1 GABA* and *GPe MEIS-SOX6 GABA*(Fig 6C), which likely correspond to the prototypic *LHX6*-expressing and arkypallidal *FOXP2*-expressing populations, respectively^73^. As in GPi, we identified homologous cell types in mouse^10,18^ for both Groups: *NDB-SI-MA-STRv Lhx8 Gaba_*4 and *GPe-SI Sox6 Cyp26b1 Gaba_1*, respectively (Fig 6C, leftmost column). In contrast to the spatial segregation of the *GPi Core* and *Shell* Groups, these GPe cell types are spatially co-mingled in all three primate species and in mouse. The proportion of prototypic to arkypallidal cells is higher in human compared to mouse, marmoset, and macaque (Fig 6C, middle row). In addition to the namesake *MEIS2*, we find that *FOXP2* is a conserved marker for the arkypallidal cell type in primates (Fig 6C bottom row, Supp Fig S14B.). Using the imputed genes from the mouse spatial transcriptomics dataset (RRID:SCR_024440), we find that both *Meis2* and *Foxp2* are also conserved marker genes in mouse (Fig. 6C, bottom row).

## Discussion

The basal ganglia are a critical component of movement, decision-making and reward functions in the mammalian brain. Because this system operates through anatomically organized circuits composed of distinct neuronal populations, spatial transcriptomic characterization of these cells is necessary to link molecular cell types to function. With more than one million cells profiled per primate species, sampled from across the full rostral-caudal extent, this spatial atlas serves as an unprecedented resource for characterizing the molecular organization of the basal ganglia nuclei. Using this resource, we find that the spatial organization of basal ganglia cell types is highly conserved across primates and mouse, suggesting that these organizational themes have functional importance preserved across millions of years of evolution. Furthermore, we identify the key principles of this conserved spatial organization. Transcriptomic cell types in the basal ganglia are organized into discrete compartments, and yet, these structures simultaneously exhibit smooth spatial variation in gene expression that spans cell types, compartments, and even anatomical boundaries. We share this spatial transcriptomic atlas via publicly available, interactive tools to enable researchers to explore additional pressing scientific questions.

### Spatial organization of the basal ganglia is conserved across primate species

Our spatial atlas demonstrates that the overall organization of cell types is highly conserved between human and non-human primates—and often all the way to mouse—despite divergences in neuroanatomy across these species. This conservation spans three aspects of spatial organization that we find in the basal ganglia nuclei: cell types that are localized to spatially segregated compartments within a nucleus (e.g., GPi and striatum), specialized cell types that are intermixed within a nucleus (e.g., GPe), and continuous gradients of gene expression that can both exist within a single nucleus (e.g., STH) and span classical nuclei boundaries (e.g., ventral vs dorsal striatum, or striatal transition regions).

We find that the GPi is organized into two discrete molecular compartments, each dominated by its own transcriptomic cell type. The primate GPi is homologous to the mouse entopeduncular nucleus (EPN), which was recently shown to have molecularly distinct core and shell divisions that are differentially innervated by striatal/pallidal sources and that route output to distinct downstream targets^71,72,75,76^. Prior axon tracing work in primates hinted at a similar spatial pattern of projections in GPi, but did not characterize the molecular identities of the cells^77–79^. Our spatial transcriptomics atlas now confirms that this core-shell organization of cell types is conserved across mouse, non-human primate, and human.

The extensive spatial resolution and genes profiled in our atlas enables us to identify conserved gene expression and cell type gradients that link the striatum to neighboring regions. This atlas identifies unique features of the amygdalostriatal transition area (AStr), a region between the caudal-ventral striatum and central nucleus of the amygdala (CEN). In mouse, AStr MSNs are more often *DRD2*-positive with distinct transcriptional profiles that appear intermediate between caudal-ventral striatum MSNs and CEN MSNs^56^. A previous study also reported spatially segregated *DRD1*-poor and *DRD2*-poor zones in the caudal-ventral striatum of mice, rats, and marmoset^80^. Our study confirms that the AStr is *DRD2*-enriched in marmoset, macaque, and human, providing further evidence for species conservation. As in mouse, *DRD2*-expressing cells in primate AStr share some features with other ventral striatum MSNs but are also transcriptionally distinct. A similar transitional pattern was observed in *DRD1*-expressing MSNs in the mediodorsal subidivsion of the nucleus accumbens shell (NACsmd) which is adjacent to the bed nucleus of the stria terminalis (BST). Taken together, these observations suggest that the AStr and NACsmd contain distinct cell subpopulations that may reflect the transitional nature of these regions and their unique functions^56,81^

### Cell types localize outside their defining neurochemical compartments

The striatum’s organization into neurochemically-defined compartments has been well-documented in both primate and mouse, and our spatial atlas characterizes compartment-associated cell types for striosomes, matrix, ICjs, and NUDAPs. However, across all primate species, we observe “exo-compartment” cells that share molecular features of their namesake compartment yet localize elsewhere. Some of these exo-compartment cells have been previously observed in the mouse, suggesting this is a conserved feature of cell type diversity and spatial organization. Here, we report a primate corollary to the mouse exo-patch cells, which resemble striosome MSNs transcriptomically but are scattered throughout the matrix compartment^40,41^. We also identify a subpopulation of primate *STRv D1 NUDAP MSNs* that are localized to the NACsmd but do not clump into canonical NUDAP compartments, a feature that has been previously noted in rat^82^. In other regions of the basal ganglia, these exo-compartment cells exhibit species-specific features. For example, in the human olfactory tubercle, we observe a much larger population of scattered granule cells outside of the Islands of Calleja, in agreement with previous studies^44^. This may have implications for odor and sensory processing differences between species: the olfactory tubercle is notably less defined in humans compared to non-human primates, and the differences are even more striking when compared to rodents, which are more smell-dependent than primates^36,83,84^. In addition to distinct molecular features, neurochemically-defined compartments often have distinct connectivity patterns. For instance, recent work showed that MSNs within striosomes project to distinct intrinsic nuclei and have opposite effects on motor function than their matrix MSN counterparts^85^. Does an exo-patch cell belong to striosome circuits or the parallel matrix circuits? It remains an open question in primates whether the connectivity and function of these exo-compartment cells resemble that of their cell type or compartment.

### Toward linking basal ganglia molecular and connectivity gradients

The relationship between gene expression and connectivity in the basal ganglia is an active area of research. In both the STH and STR, our spatial atlas uncovers gene expression gradients that are strikingly similar to published connectivity gradients, suggesting a potential correspondence. Injection tracing studies have revealed that corticostriatal projections are generally characterized by diffuse, overlapping projections from specific regions of frontal cortex, organized along a ventromedial-to-dorsolateral gradient^20^. A parallel topographic organization has been identified in the STH, an important clinical target for deep brain stimulation (DBS) in the treatment of motor symptoms of Parkinson’s Disease^86,87^, with overlapping functional domains along its longitudinal axis^57^. Related investigations in human have used fMRI approaches and confirm that connectivity in STR and STH includes highly overlapping projection fields^58,88^, though other studies emphasize more discrete organization^89,90^.

Here, we find a variety of smooth gene expression patterns within the dominant gradients of the striatum, but it remains unclear if connectivity is correlated to a specific gene, to the overall gradient, or to another organizing principle. Also missing is a link between the anatomical localization and the functional roles of the genes measured in this study. For example, MSNs in STRd exhibit a gradient of expression of *CNR1*, the gene for the CB1 cannabinoid receptor, which is known to modulate synaptic transmission presynaptically at MSN projection targets^91^ and has a dorso-ventral gradient of ligand binding *in vivo*^92^. Do individual neurons projecting from cortex innervate MSNs with a narrow or a wide range of *CNR1* expression? Given *CNR1*’s role in neuromodulation, this link between gene expression and connectivity has direct relevance to basal ganglia circuit function.

Multimodal experiments will be needed to address these gaps in our understanding of basal ganglia organization. One such experiment could measure both connectivity and spatial transcriptomics in the same cells to link individual transcriptomic cell types to connectivity properties, perhaps using high-throughput methods such as BARseq^93^ or new labeling and imaging approaches for large brains^94^. Another approach could compare detailed fMRI measurements with spatial transcriptomics measurements, following registration of our atlas to MRI volumes. In both of these approaches, this spatial transcriptomic atlas can provide a common framework for the future multi-modal datasets that will be crucial for linking striatal cell types to functional connectivity.

### Limitations

Spatial transcriptomics is an emerging technology with experimental and analysis techniques that are still maturing. The detection efficiency of mRNA transcripts varies based on the underlying chemistry and imaging used in different commercial platforms, which are not yet accounted for in standard sc/snRNA-seq approaches for gene count normalization and transformations^95^. Cell segmentation of spatial transcriptomics data is another active area of development. While we found that the on-instrument Xenium segmentation staining kit and associated algorithm offered a good balance of simplicity and accuracy, the on-instrument MERSCOPE segmentation was prone to significant error; thus, we co-developed a more robust cell segmentation pipeline utilizing Cellpose 2.0 human-in-the-loop and training across multiple species^96^. We saw visible and quantitative improvement in segmentation results using this pipeline on our macaque and human MERSCOPE data.

Spatial transcriptomic cells are often assigned an identity through label transfer (mapping) methods which are designed to map unlabeled sc/snRNA-seq data to an annotated sc/snRNA-seq reference dataset. Spatial transcriptomics mapping results can be imperfect due to gaps in the reference data, technical differences between spatial and RNA-seq transcript measurements, the gene panel, and the computational approach for mapping. Using current methods, mapping ambiguity most often impacts cells with closely related transcriptomic identities. For instance, the *STRv D1 NUDAP MSNs* and *STR D1D2 Hybrid MSNs* Groups are transcriptomically similar and are known to exist within a larger continuum of “eccentric” MSN types^97^. In our spatial transcriptomic atlas, this may manifest as colocalization or ambiguous mapping. The same phenomenon could impact other closely related cell types, such as the exo-patch cells that may be an intermediate between the striosome and matrix MSNs. When interpreting spatial localization of highly related clusters, identifying spatially variable genes can help to verify the results. Despite these considerations, the high correlation of Group populations across modalities and species bolsters the utility of this atlas. Cell type mapping and multimodal integration is an actively developing field^98,99^, and future spatial atlases will benefit from ongoing efforts to optimize these algorithms.

Finally, the larger size of primate brains also poses challenges for spatial transcriptomic profiling. Coronal sections of an entire marmoset hemisphere fit within the Xenium imaging area, while coronal tissue slabs from human and macaque had to be subdivided into smaller blocks to fit in the MERSCOPE imaging area (Fig S14A). Although this posed significant challenges, careful alignment and registration (*Methods*) enabled us to place adjacent sections into common anatomical coordinates, leaving only minor gaps and discontinuities between sections (Fig S14B). Future spatial transcriptomics platforms may enable profiling of larger sections and reduce the need for registration.

### Toward whole-brain primate atlases

The BRAIN Initiative Cell Atlas Network (BICAN) is a collaboration among neuroscientists to create comprehensive, multi-modal, whole-brain atlases of the non-human primate and human brains. On its own, this basal ganglia spatial transcriptomics atlas makes significant progress in characterizing the cell type and molecular organization of basal ganglia nuclei. However, many scientific questions will only be answered by profiling the entire brain. Many cells captured outside of our anatomically defined basal ganglia structures either map to cell types beyond the consensus taxonomy or reveal that these types may have counterparts in other structures.

Further, we identified several transition regions that are conserved across species and are often overlooked in single region studies. The multi-modal, cross-species basal ganglia atlas presented here and in companion papers (https://brain-map.org/consortia/hmba/hmba-release-basal-ganglia) represents an important foundational step toward the whole-brain atlases to come.

## Acknowledgements

We thank Song-Lin Ding for conversations on anatomical delineations, the Basal Ganglia Analysis Working Group for discussions around cell typing, and Chelsea Pagan for program management support. We also thank Cliff Slaughterbeck and the SIPE team for instrumentation and software support. A thank you to Raymond Sanchez, Elysha Fiabane, Chris Morrison, Scott Daniel, and Tyler Mollenkopf of the Data and Technology team for product development. We thank the animal care personnel and veterinary staff at MIT DCM and Rockefeller for their dedicated help with animal husbandry and expert clinical support. The authors thank the Allen Institute founder Paul G. Allen for his vision, encouragement and support, and Allen Institute chair Jody Allen. This publication was supported by and coordinated through the Brain Initiative Cell Atlas Network (BICAN). Research reported in this publication was funded by the National Institute of Mental Health under NIH award UM1MH130981-01 and the Allen Institute for Brain Science.

## Materials and Methods

### Tissue procurement/sourcing Human

Donors 16 – 68 years of age with no known history of neuropsychiatric or neurological conditions (‘control’ cases) were considered for inclusion in this study. De-identified postmortem human brain tissue was collected after obtaining permission from the decedent’s legal next-of-kin. Tissue collection was performed in accordance with the provisions of the United States Uniform Anatomical Gift Act of 2006 described in the California Health and Safety Code section 7150 (effective 1/1/2008) and other applicable state and federal laws and regulations. The Western Institutional Review Board (WIRB) reviewed the use of de-identified postmortem brain tissue for research purposes and determined that, in accordance with federal regulation 45 CFR 46 and associated guidance, the use of de-identified specimens from deceased individuals did not constitute human subjects research requiring IRB review. Routine serological screening for infectious disease (HIV, Hepatitis B, and Hepatitis C) was conducted using donor blood samples and donors negative for all three infectious diseases were considered for inclusion in the study. Tissue RNA quality was assessed using samples of total RNA derived from the frontal and occipital poles, which were processed on an Agilent 2100 Bioanalyzer using the RNA 6000 Nano kit to generate RNA Integrity Number (RIN) scores for each sample. The donor tissue featured in this study originated from a 50-year-old Caucasian female who died of natural causes, with a postmortem interval of 8.1 hours. RIN values for this tissue upon intake were ≥8.0 for all regions assessed. At the point of block creation and data collection, RIN values specifically taken for the basal ganglia regions profiled were ≥8.0. The right hemisphere was profiled for spatial transcriptomics.

### Macaque

The brain of one male Rhesus macaque monkey (HMBA ID: QM23.50.001) aged 10 years was collected for use in this study and the right hemisphere was profiled for spatial transcriptomics. All procedures for brain extraction were performed in The Rockefeller University’s AAALAC accredited surgical suite under IACUC approved protocol# 24066-H and in compliance with all applicable federal and state animal welfare laws, regulations, and policy. In order to maximize the quality of the tissue to be subjected to gene-expression analysis, we attempted to minimize the time between the beginning of the transcardial perfusion and freezing. After initial sedation in the home cage using Ketamine (∼3-10 mg/kg) plus Dexdomitor (∼0.001-0.02 mg/kg) administered IM, the subject was intubated, and intravenous catheters were placed in the surgical suite. The subject was placed on a surgical table with heat support provided by a water recirculating blanket and a forced-air warming system, life support provided by a mechanical ventilator based on animal ETCO2 levels and maintained on intravenous lactated ringers throughout the duration of anesthetized surgical procedures, and vital monitoring including respiratory rate, heart rate, blood pressure, end tidal CO2, Sp02 and core body temperature, and its head was placed in a stereotaxic frame. Once all vital parameters were stable, we established a plane of deep level of anesthesia using continuous infusions of intravenous fentanyl (∼3-25 mcg/kg/hr) and dexdomitor (∼1-3 mcg/kg/hr) along with gas isoflurane (∼0.25-2.5%). Level of anesthesia was confirmed using heart rate and toe pinch response as well as respiratory rate and palpebral response. Vitals and anesthesia levels were independently monitored by a veterinary technician and a veterinarian. Once this deep level of anesthesia was achieved, the skull was exposed and opened, and the dura mater removed. From this point on the brain was continuously flooded with a physiological saline solution to prevent it from drying. Upon confirmation of the depth of anesthesia by the veterinarian, transcardial perfusion was prepared by IV injection of pentobarbital sodium and phenytoin sodium at >150mg/kg to induce cardiac arrest. When this was confirmed by the veterinarian, an aortal catheter was introduced and transcardial perfusion with PBS 1X initiated, which lasted for ten minutes. Subsequently, the brain was removed in about two minutes and placed into a mold for slabbing (see below).

### Marmoset

The brain of one five-year-old male marmoset (MIT ID 18-109, HMBA ID CJ23.56.004) was used in this study and the right hemisphere was profiled for spatial transcriptomics. All marmoset experiments were approved and conducted in compliance with the Massachusetts Institute of Technology CAC (IACUC) under protocol number 2303000479. The animal was initially sedated with Alfaxalone (12mg/kg, 10 mg/ml) and Midazolam (0.3 mg/kg, 5mg/ml) via intramuscular injection. They were further sedated with a secondary dose of Alfaxalone (4mg/kg, 10mg/ml) to ensure deep sedation followed by an intravenous injection of Euthasol (>120mg/kg, 390 mg/ml). When respiration and the pedal withdrawal reflex were eliminated, the marmosets were transcardially perfused with ice-cold carbogenated N-methyl-D-glucamine (NMDG) artificial cerebrospinal fluid (aCSF) (92 mM NMDG, 2.5 mM KCl, 1.25 mM NaH₂PO₄, 30 mM NaHCO₃, 20 mM HEPES, 25 mM glucose, 2 mM thiourea, 5 mM sodium L-ascorbate, 3 mM sodium pyruvate, 0.5 mM CaCl₂·2H₂O, and 10 mM MgSO₄·7H₂O; pH 7.3-7.4 adjusted with HCl). The brain was extracted from the skull and placed in ice-cold sucrose-HEPES. The brainstem and cerebellum were cut off and the brain was placed in a chilled brain mold and sliced with a razor blade into 5 mm slabs.

### Sampling plans

#### Slabbing

Fresh marmoset, macaque, and human hemispheres were slabbed in the coronal plane following the post-mortem brain processing procedure (https://dx.doi.org/10.17504/protocols.io.bf4ajqse). Marmoset tissue was slabbed at ∼5 mm thickness resulting in 6 slabs from rostral to caudal, macaque tissue was slabbed at ∼5-7 mm thickness resulting in 12 slabs, and human tissue was slabbed at ∼4 mm thickness resulting in 60 slabs. Slabs were flash frozen, vacuum sealed, and stored in -80°C until needed. Fresh and frozen slabs were photodocumented for downstream data alignment.

#### Blocking

Selected human and macaque slabs were equilibrated to -20°C for 1 hour prior to blocking (Supp Fig S15A). Areas of interest within the slab were identified, outlined, and divided into blocks with a surface area smaller than that of the imaging area designated by the MERSCOPE platform. Blocking was performed while maintaining slabs and blocks at -20 C for the duration of the procedure. Each block and its relation to other blocks were photodocumented to ensure proper transformation of resulting data relative to other blocks and the slab as a whole.

#### Sectioning

Frozen tissue blocks were affixed to a sectioning chuck using Optimum Cutting Temperature medium (VWR 25608-930) and sectioned on a Leica cryostat at -17C at 10 μm onto Vizgen MERSCOPE coverslips (macaque and human) or 10X Xenium slides (marmoset). Marmoset hemispheres were kept intact for sectioning as they fit within the imaging window of 10X Xenium slides (Supp Fig S15C). Sections from selected slabs were collected at 1 mm spacing intervals for macaque and human and at 200 µm intervals for marmoset (Fig 1 A-C). The sampling distance between slabs is variable and undefined due to a small amount of tissue loss at the beginning and end of the slab. Photos of the block face were taken prior to section collection for downstream data registration. Back-up sections for each primary one were collected and stored according to the platform specifications.

#### Gene Panel Design

Gene panels of up to 300 genes were designed for species (300 each in marmoset and macaque, 299 in human) and spatial platform specifically with an effort to select overlapping genes when possible; 72 genes are shared across the three species gene panels (Table S1). The macaque and human MERSCOPE panels were designed in similar ways using a combination of tools. First, marker genes for the basal ganglia were chosen by hand with an effort toward selection of conserved genes across species. These were used as a starter list for two gene selection tools, mFISHtools^100^ and geneBasis^101^. mFISHtools selects marker genes in a label-aware fashion with a supplied taxonomy, in this case an early version of the consensus taxonomy. In contrast, geneBasis selects genes label-free and instead tries to maximize the distance between a low-dimension manifold of reference RNA-seq data with all available genes and the same representation with the reduced gene selection. In this way, geneBasis can suggest genes that capture expression variance that is not captured in cell types, and we find using this tool in combination with mFISHtools to be a useful strategy. The marmoset Xenium gene panel was similarly constructed as a combination of computationally and manually selected genes. The initial manual genes were selected from published marker genes, genes shared between human and macaque panels, and genes showing differential expression between Caudate and Putamen structures in the RIKEN ISH atlas (“Differential Search” tab at https://gene-atlas.brainminds.jp/gene-structural/). These genes served as starting genes for further computational search with GeneBasis and mFISHtools using data from a previous study^27^.

After candidate gene panels for each species were designed, they were reviewed by 10X or Vizgen for compatibility with the relevant platform. Certain genes (e.g., *PVALB*, *HTR2C*) were replaced for not meeting the technical requirements for making probes such as transcript length, number of target sites, and expression level. In the case of the marmoset panel, iterations with the 10X probe design tool (https://www.10xgenomics.com/support/software/xenium-panel-designer/) also optimized the panel for utilization across cell types.

### MERSCOPE methods

Sections were allowed to adhere to Vizgen MERSCOPE coverslips at room temperature for 10 minutes prior to a 1 minute wash in nuclease-free phosphate buffered saline (PBS) and fixation for 15 minutes in 4% paraformaldehyde in PBS. Fixation was followed by 3x5 minute washes in PBS prior to a 1 minute wash in 70% ethanol. Fixed sections were then stored in 70% ethanol at 4C prior to use and for up to one month. Human sections were photobleached using a 240W LED array for 72 hours at 4°C (with temperature monitoring to keep samples below 17°C) prior to hybridization then washed in 5 mL Sample Prep Wash Buffer (VIZGEN 20300001) in a 5 cm petri dish. Sections were then incubated in 5 mL Formamide Wash Buffer (VIZGEN 20300002) at 37°C for 30 min. Sections were hybridized by placing 50 μL of VIZGEN-supplied Gene Panel Mix onto the section, covering with parafilm, and incubating at 37°C for 36-48 hours in a humidified hybridization oven. Following hybridization, sections were washed twice in 5 mL Formamide Wash Buffer for 30 minutes at 47°C. Sections were then embedded in acrylamide by polymerizing VIZGEN Embedding Premix (VIZGEN 20300004) according to the manufacturer’s instructions. Sections were embedded by inverting sections onto 110 μL of Embedding Premix and 10% Ammonium Persulfate (Sigma A3678) and TEMED (BioRad 161-0800) solution applied to a Gel Slick (Lonza 50640) treated 2x3 inch glass slide. The coverslips were pressed gently onto the acrylamide solution and allowed to polymerize for 1.5 hours.

Following embedding, sections were cleared for 24-48 hours with a mixture of VIZGEN Clearing Solution (VIZGEN 20300003) and Proteinase K (New England Biolabs P8107S) according to the manufacturer’s instructions. Following clearing, sections were washed 2x5 minutes in Sample Prep Wash Buffer (PN 20300001). VIZGEN DAPI and PolyT Stain (PN 20300021) was applied to each section for 15 minutes followed by a 10 minutes wash in Formamide Wash Buffer. Formamide Wash Buffer was removed and replaced with Sample Prep Wash Buffer during MERSCOPE set up. 100 μL of RNAse Inhibitor (New England BioLabs M0314L) was added to 250 μL of Imaging Buffer Activator (PN 203000015) and this mixture was added via the cartridge activation port to a pre-thawed and mixed MERSCOPE Imaging cartridge (VIZGEN PN1040004). 15 mL mineral oil (Millipore-Sigma m5904-6X500ML) was added to the activation port and the MERSCOPE fluidics system was primed according to VIZGEN instructions. The flow chamber was assembled with the hybridized and cleared section coverslip according to VIZGEN specifications and the imaging session was initiated after collection of a 10X mosaic DAPI image and selection of the imaging area. Specimens were imaged and automatically decoded into transcript location data.

### Xenium methods

Fresh frozen tissue sections were mounted onto Xenium slides (10X Genomics) and stored at −80 °C until use. Slides were equilibrated at 37 °C for 1 min using a pre-heated thermal cycler (Bio-Rad 1851197) equipped with a Xenium Thermocycler Adaptor. Fixation was performed in 1× PBS containing 2.5 mL paraformaldehyde (Electron Microscopy Sciences 15710) for 30 min at room temperature. Slides were permeabilized with 1% SDS (Millipore Sigma 71736) and incubated in pre-chilled 70% methanol for 60 min on wet ice.

Following permeabilization, slides were washed and transferred into Xenium cassettes. Probe hybridization was performed using a mix of Xenium Probe Hybridization Buffer, Probe Dilution Buffer, and either pre-designed or custom gene expression probes (10X Genomics). Probes were preheated at 95°C for 2 min and cooled on ice prior to mixing. Each slide received 500 µL of hybridization mix and was incubated overnight (16–24 h) at 50°C. Following hybridization, slides underwent washes using Xenium Post Hybridization Wash Buffer at 37°C. Ligation was performed by adding 500 µL of freshly prepared ligation mix containing Xenium Ligation Buffer, Enzyme A, and Enzyme B, followed by a 2 h incubation at 37°C. Amplification was conducted using a master mix of Xenium Amplification Mix and Enzyme, incubated for 2 h at 30°C. Autofluorescence quenching was achieved by sequential washes in TE buffer, 1x PBS, and ethanol solutions, followed by incubation with Xenium Autofluorescence Solution for 10 min in the dark. Nuclei staining was performed using Xenium Nuclei Staining Buffer, followed by four washes in PBS-T. Slides were stored in PBS-T at 4 °C or immediately processed for imaging.

Prepared slides were loaded into the Xenium Analyzer (10X Genomics 1000481) along with decoding reagent modules, buffer bottles, and consumables. Imaging buffers were freshly prepared, including Xenium Probe Removal Buffer and Sample Wash Buffers A and B. Reagents were loaded into designated positions on the instrument, and system checks were performed prior to initiating the run. Following the run, slides were scanned for DAPI and autofluorescence signals, and imaging regions were designated using the instrument’s touchscreen interface. Imaging runs were initiated and completed over 1–3 days. Upon completion, fluidics cleanup was performed, and slides were stored in PBS-T at 4 °C. Imaging data were exported and reviewed for quality control.

### Section and Block Alignment

The blocking procedure utilized for macaque and human tissue necessitated registration and stitching of sections, along with the spatial transcriptomics data, back to the original slab tissue coordinates (Supp Fig S15B). Spatial transcriptomic sections were first registered to their corresponding block-face images using affine transformations. Visual inspection confirmed that tissue deformations in the imaged sections were minimal relative to the block-face images acquired immediately prior to sectioning. Block-face images were subsequently registered to slab tissue images to anchor the sections in slab space. The resulting stitched spatial transcriptomics image thus represents the same sectioning plane across independent blocks and was treated as a continuous coronal plane in downstream analysis (Supp Fig S15B, far left). Prior to registration, image scales were normalized and orientations were standardized to neurological convention (right hemisphere displayed on the right). Tissue cut surfaces were segmented and cropped to isolate the relevant block-face and slab-face regions, removing background and extraneous tissue outside the section plane. Anatomical landmarks were manually identified to perform affine registration between the block-face section most closely corresponding to each slab image. Finally, block-face images across consecutive sections within each block were aligned by matching cut edges and ensuring continuity of anatomical features between adjacent mosaicked blocks.

### Spatial Data Post-Processing

Post-processing of the MERSCOPE and Xenium data was conducted utilizing custom tools with a focus on consistency and reproducibility across the three datasets (Supp Fig S2). While the MERSCOPE platform offers cell segmentation tools, we chose to re-segment macaque and human data with our custom segmentation pipeline that shows improved data quality. The software is available on Github (https://github.com/AllenInstitute/spots-in-space/tree/main/sis). In brief, re-segmentation of the macaque and human MERSCOPE data was performed using a custom 3D Cellpose2.0^32^ model that was trained on mouse, macaque, and human MERSCOPE data using the human-in-the-loop approach. The model utilized two channels, the DAPI image acquired during MERSCOPE acquisition and an image created from the total mRNA signal of the MERSCOPE data to serve as a cytosol stain (Supp Fig S2A). Cell segmentation of marmoset data was performed on the Xenium instrument using the segmentation kit. Cells were deemed of low quality if they failed to exceed thresholds on the total number of transcripts (marmoset: 20, macaque: 20, human: 10) and the number of unique genes detected in each cell (marmoset: 3, macaque: 6, human: 4) (Supp Fig S2B). Additionally, the MERSCOPE gene panels include “blank” codewords that are not assigned to any real gene probe in the panel.

These assess decoding error and can be utilized as a floor to the robustness of gene decoding. An upper limit of 2% of transcripts in a cell being assigned to a “blank” codeword was also applied to the macaque and human data. All cells are included in the data release but cells that do not meet the above QC thresholds are excluded from further analysis using the ‘qc_pass’ flag (Supp Fig S2D).

### Manual Anatomical Annotations

To place the spatial transcriptomics data into the context of anatomical regions manual outlines for the five major regions of the basal ganglia (STR, GP, STH, SN) were drawn onto every section for all three species (Supp Fig S1). These were chosen for their consistency and definitiveness throughout the data. Though these do not necessarily align anatomically with the corresponding regions in the Harmonized Ontology of Mammalian Brain Anatomy (HOMBA)^31^, the same acronyms are used for consistency. Additionally, in keeping with HOMBA ontology we refer to the longitudinal axis (front to back) of these structures as rostral (R) to caudal (C).

Manual anatomical annotations were drawn using the polygon tool in napari^102^. For the marmoset data, the Xenium cell segmentation kit stains were used to identify boundaries between white and grey matter. Further anatomical delineations were drawn using a combination of stains and marker gene expression (Supp Fig S1A). The marmoset Paxinos atlas was used as a reference^103^. For the macaque and human MERSCOPE data, which did not include additional stains, annotations were drawn using a combination of marker gene expression (Supp Fig S1B-C, Supp Table S8), cell density, and preliminary spatial domains (see “Spatial domain detection” below). The macaque Paxinos atlas^104^ and the Allen Human Brain Atlas^105^ were used as references. Finally, shapely’s “contains” function was used to label the cells that fell inside each of the manual annotation polygons drawn in napari. As the manually drawn polygons sometimes overlapped, cells were allowed to belong to multiple anatomical annotations.

### Spatial domain detection

For the macaque and human datasets, domain detection was performed on binned transcript locations using STAligner^106^. Binned transcripts were used rather than segmented cells to account for GPU memory limitations. For macaque, transcripts were assigned to 50 micron square bins and bins with fewer than 60 unique genes were filtered out. For human, transcripts were assigned to 50 micron square bins and bins with fewer than 30 unique genes were filtered out. STAligner embedding was run on all sections from a species, with each section being considered a separate subgraph for training. Following the embedding, clustering was performed on the aligned latent space. For macaque, the leiden algorithm was used with resolution 0.8 and 15 nearest neighbors. For human, the mclust algorithm was used to identify 10 clusters.

### Analysis approach

#### Celltype mapping

Cells from each species were mapped to their respective snRNA-seq reference datasets using MapMyCells’ (RRID:SCR_024672) (Supp Fig S2C) flat mapping algorithm with bootstrapping (100 iterations, bootstrap factor 0.95). For macaque and human, the snRNA-seq preprint taxonomy with Adjacent clusters was used as a reference. Cluster Human-382 was removed from the human reference before mapping. For marmoset, a full subcortex snRNA-seq taxonomy published concurrently^107^ with v3 clusters was used as a reference. Using the bootstrapping probabilities, the Shannon entropy was calculated at the Group level for each cell, and cells with Group Shannon entropy > 0.7 were filtered out before analysis.

#### Analysis details (packages, software etc.)

Data used for further analysis included only those cells that passed cell-segmentation QC (‘qc_pass’), was within the boundaries of the manually annotated basal_ganglia_and_adjacent region, and mapped robustly (Group Shannon entropy > 0.7) to the Basal Ganglia Consensus Taxonomy. Data was organized from most rostral to most caudal and is presented in figures as such (ex. Fig 1A-C). Unless otherwise specified, gene expression data for macaque and human are presented as log2CPT (counts per thousand) and as raw counts for marmoset. These differing normalization approaches reflect the difference in detection sensitivity between the spatial transcriptomics platforms.

#### Compartment masks

Polygon masks for the three compartments in the striatum were generated from the cells belonging to their associated Groups: striosomes (*STRd D1 Striosome MSN*, *STRd D2 Striosome MSN*), ICjs (*OT D1 ICj*), and NUDAPs (*STRv D1 NUDAP MSN*, *STR D1D2 Hybrid MSN*). For each compartment, cells with the Group labels were initially filtered to remove extreme outliers first using a KNN with 5 nearest neighbors and then using scipy’s DBSCAN algorithm (minimum cluster size, all species: 10). Remaining cells were binned (marmoset: 20 µm; macaque: 50 µm; human: 80 µm) and converted to a binary mask. After applying “binary_closing” and “binary_fill_holes” functions, the binary masks were converted first to contours using the cv2 package and finally shapely polygons. Individual polygons were given unique identifiers and cells were assigned to the specific polygon they fell inside via shapely’s contains function.

#### Principal component analysis (PCA)

Principal component analysis (PCA) was used to analyze gene expression gradients. Prior to PCA, cells were subset to the Groups of interest (for STRd, *STRd D1 Matrix MSN*, *STRd D2 Matrix MSN*, *STRd D1 Striosome MSN*, and *STRd D2 Striosome MSN*; for STRv, the Groups used for STRd plus *STRv D1 MSN*, *STRv D2 MSN*, and *AMY-SLEA-BNST GABA*). For rostral STRd and STRv (Fig 3 and Supp Fig S9; Fig 4, top half), a manual line was drawn to indicate the overall direction of the internal capsule. For caudal STRv, the line was drawn from the central nucleus of the amygdala (inferred from the presence of *AMY-SLEA-BNST GABA* cells) to the lateral edge of the caudate tail (Fig 4, bottom half). In both cases, cell coordinates were projected onto the manually drawn axis, and PCA (scikit-learn) was performed on each species separately to maximize the number of genes. The principal gradient components (PGCd and PGCv were chosen as the PCs with highest correlation to the cells’ projected coordinates.

PGCd direction shown in Figure 3 (green arrow in top panel) is measured from a 2D linear fit to the PGCd values in cells of each Group.

#### Mouse Data

Mouse spatial transcriptomics data is part of the Allen whole mouse brain cell type atlas^10^ and is available for download through the ABC Atlas (RRID:SCR_024440). Datasets MERFISH-C57BL6J-638850 with Imputed Genes + Reconstructed Coordinates and MERFISH-C57BL6J-638850 Reconstructed Coordinates were used for comparison with primate species in the GP (Fig 6).

#### Cytosplore Gradient Surfer

Gradient Surfer is a plugin for the Cytosplore Viewer data visualization system^37^, available at https://viewer.cytosplore.org including spatial data from this paper. Starting with one dataset as reference, the user can interactively select cells from the data, e.g., cells of the dorsal striatum, in a linear strip along the internal capsule axis. Cell positions of selected cells are projected on the line, and Pearson correlation is calculated between the expression profiles of the shared gene set and the projected cell coordinates. This reveals genes whose expression consistently varies with or against the line coordinate, reflecting a gradient along the drawn line. Principal Component Analysis (PCA) is applied to the matrix of the selected cells by the top N gradient genes, resulting in a set of gradient PC’s for the selected cells in the reference dataset. The PC with the strongest correlation with the linear position along the line is automatically selected.

The second dataset may be acquired with a different gene panel, but no line is selected, so a second PCA is performed on all cells in the second dataset, but only on the top N gradient genes as calculated in the reference dataset. The PC with the strongest correlation of the reference gradient PC is selected as the best match. Within each dataset, all genes are ordered by correlation with the strongest (or manually selected) gradient PC for that dataset. However, PC loadings can be visualized as color overlays on spatial scatterplots similar to the gene expression to verify which PC represents the probed gradient best. See Supplemental Figure 7.

