## Supplemental Information for "A cross-species spatial transcriptomic atlas of the human and non-human primate basal ganglia"

**Supplemental Table S1:** Full gene list across three species spatial transcriptomics. Each column identifies if that gene is present in that species

**Supplemental Table S2:** Consensus taxonomy hierarchy and Group short names.

**Supplemental Table S3:** Cell count and proportion for each Group in each of the manually annotated regions in marmoset.

**Supplemental Table S4:** Cell count and proportion for each Group in each of the manually annotated regions in macaque.

**Supplemental Table S5:** Cell count and proportion for each Group in each of the manually annotated regions in human.

**Supplemental Table S6:** Neuron Group proportions across species and modalities.

**Supplemental Table S7**: Consolidated regional proportions for each Group across species with average.

**Supplemental Table S8:** Genes used to aid in manual anatomical annotation drawings.

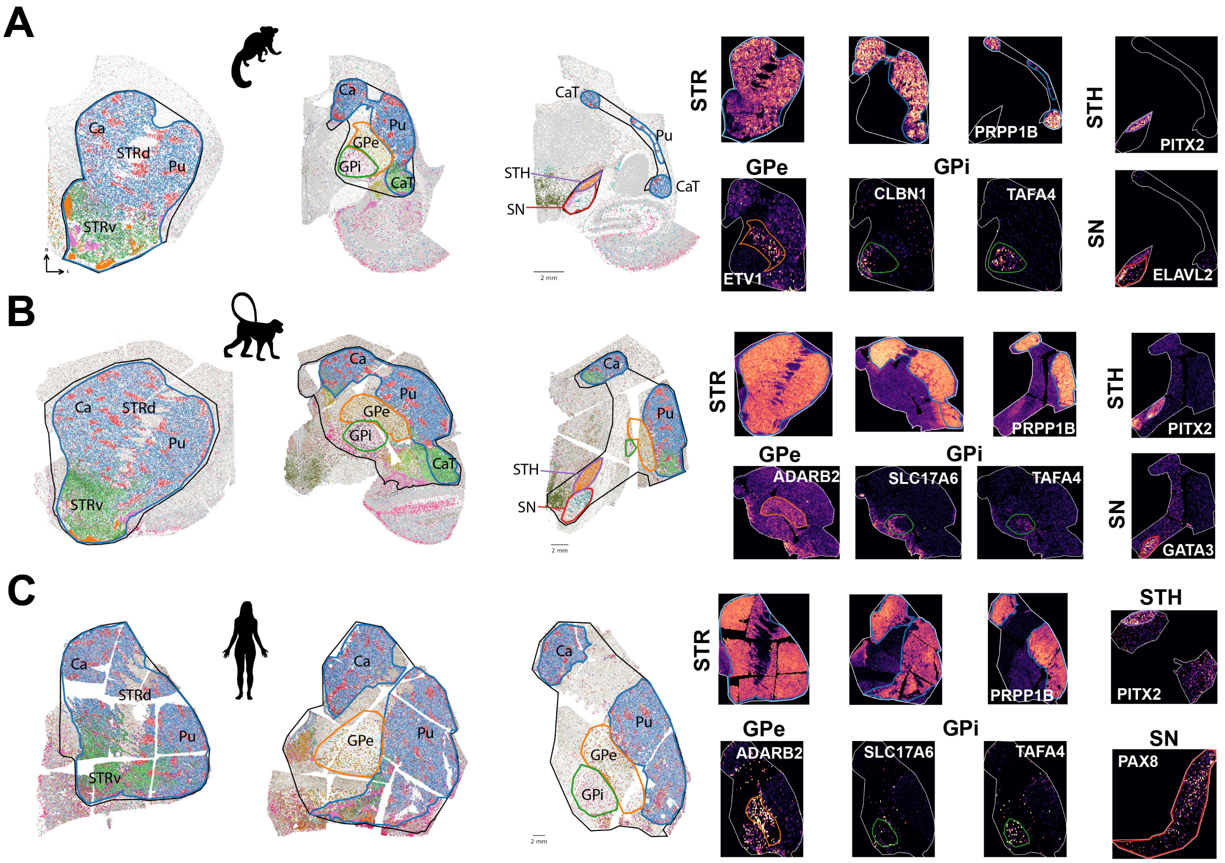


**Supplemental Figure S1:** Manual anatomical annotations, related to Figure 1

For each species (A-C) manual outlines were drawn for anatomical regions including the striatum (STR, blue), GPe (orange), GPi (green), subthalamic nucleus (STH, purple), and substantia nigra (SN, red). While they could not be reliabely outlined, the caudate (Ca), caudate tail (CaT), and putamen (Pu) are identified as landmarks. These regions were encompassed in a larger outline (Adj, black) for completeness. Left: slices from each species at three different rostral to caudal positions, with cells colored by Group and each of these annotations outlined. Right: histogram expression of individual genes used to guide the drawings. When possible, similar genes were used across species (Supp Table S7). The Adj outline in white is included for orientation.

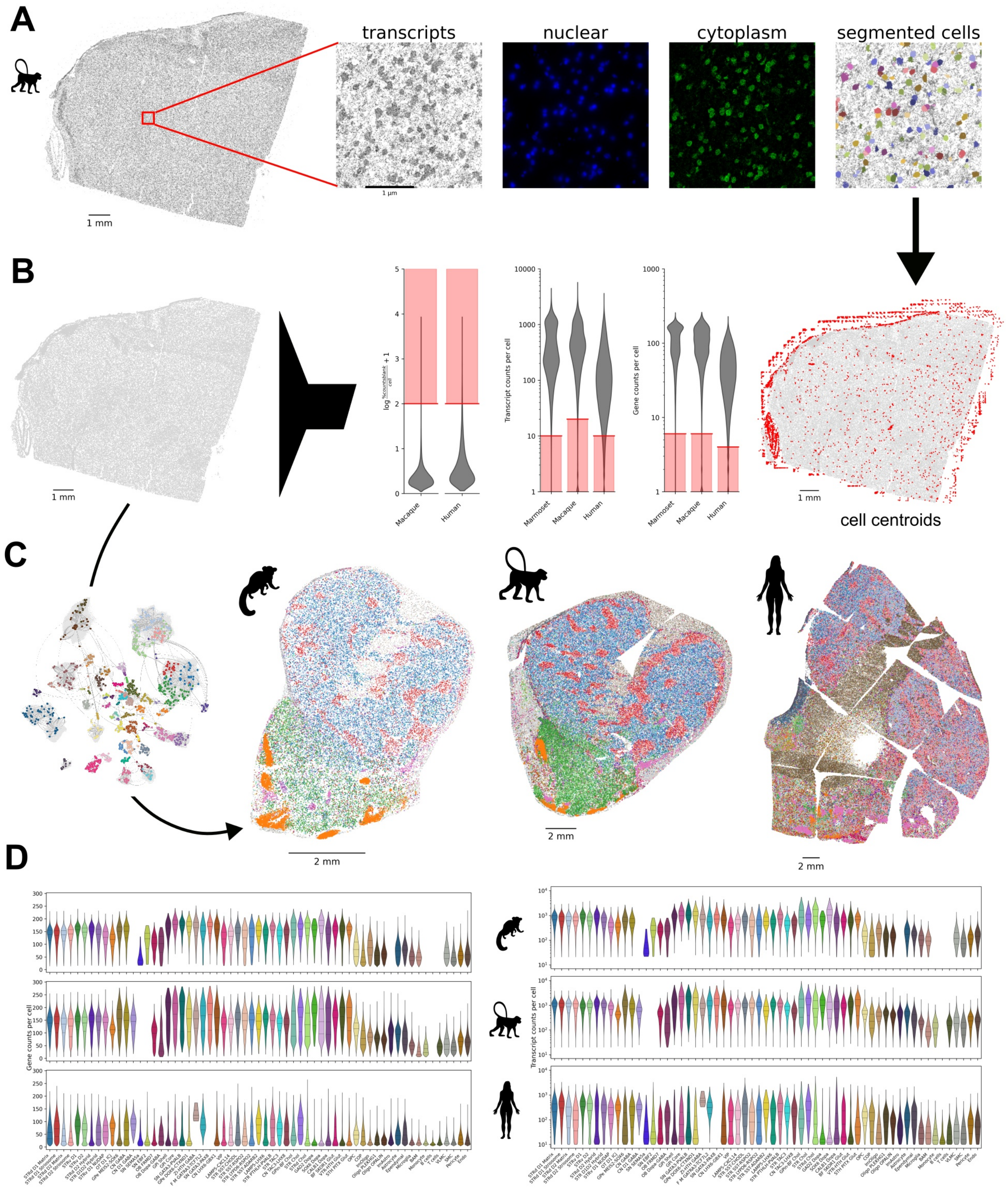


**Supplemental Figure S2:** Post-processing of spatial transcriptomics data, related to Figure 1, Methods

A and B use an example macaque section to depict the spatial transcriptomics data processing workflow that was applied to data from all species. **A.** Cell segmentation of spatial transcriptomics sections utilized varying combinations of transcript density, nuclear, and cytoplasmic stains depending on the acquisition platform (see Methods). **B.** Segmented cells were QC'ed using thresholded parameters including number of transcripts and genes per cell. **C.** QC'ed cells were mapped to the BG consensus taxonomy using MapMyCells, here showing an example slice of striatum from each species colored by Group. **D.** Distributions of number of transcripts and number of genes detected per cell after QC and mapped to each Group for each species. Violins colored by Group and inner lines depict median (large dash) and quartiles (small dash). Shortened Group names are used (see Supp Table S2).

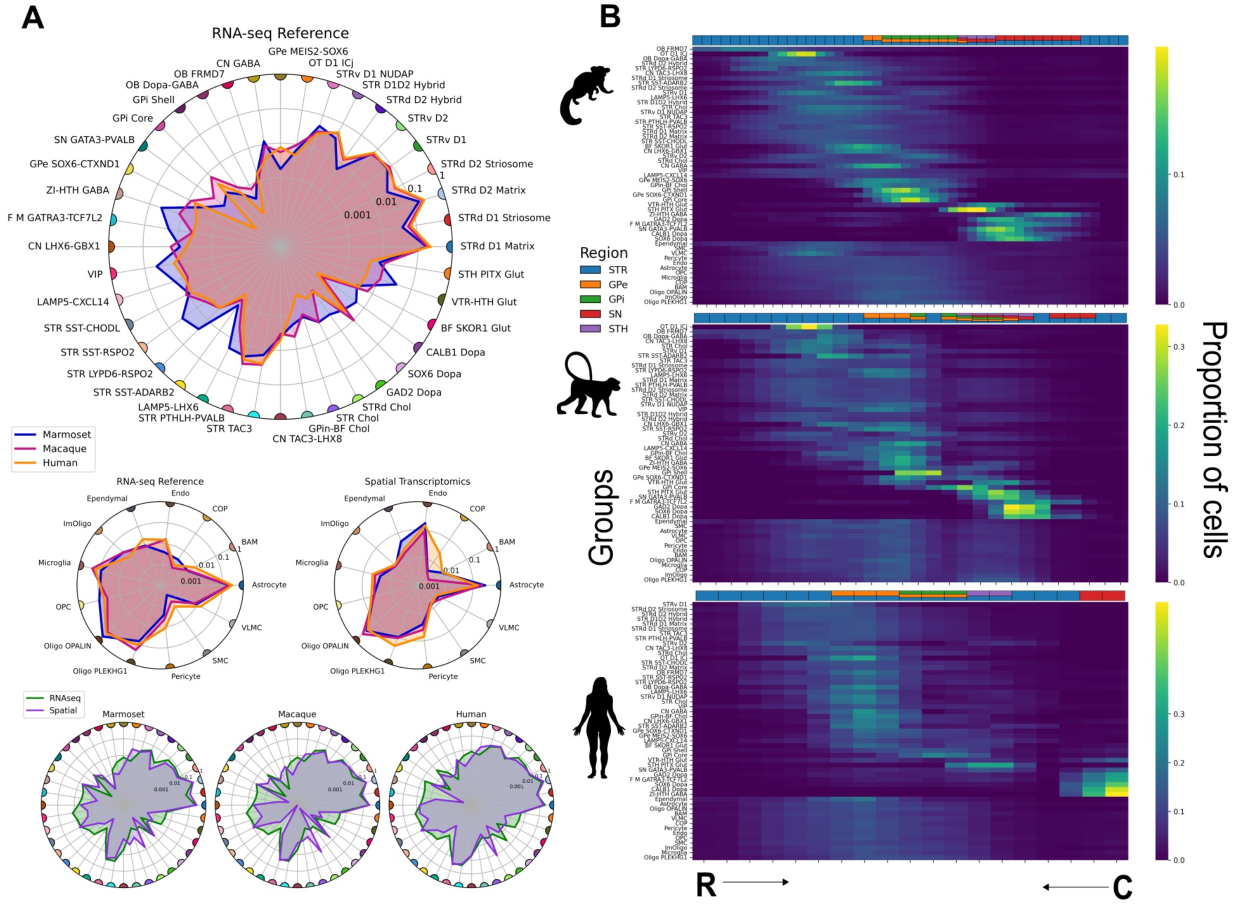


**Supplemental Figure S3:** Group proportions related to snRNAseq and R-C axis, related to Figure 1

**A.** Neuron Group (short names, Supp Table S2) proportions in each species in RNAseq reference data (top). Non-neuron Group proportions in snRNAseq (left) and spatial transcriptomics (right) for each species (middle). Comparison of neuron Group proportions across modalities in each species. Colored dots at each spoke follow from the top plot. For all radial proportion plots the denominator consists only of the Groups depicted in the plot (ie the sum is 1). **B.** Heatmap of Group (short names) proportions across sections in the rostral (R) to caudal (C) axis. For each Group (row) a moving average was computed over 5 sections in marmoset and 3 sections in macaque and human. Groups are sorted with neuron Groups on top and non-neuron Groups on the bottom. With in these two, for each species, Groups are sorted by the center of mass (CoM) in the rostral-caudal axis. Each row sums to 1. Above each heatmap is a stacked bar chart depicting the region(s) present in each section.

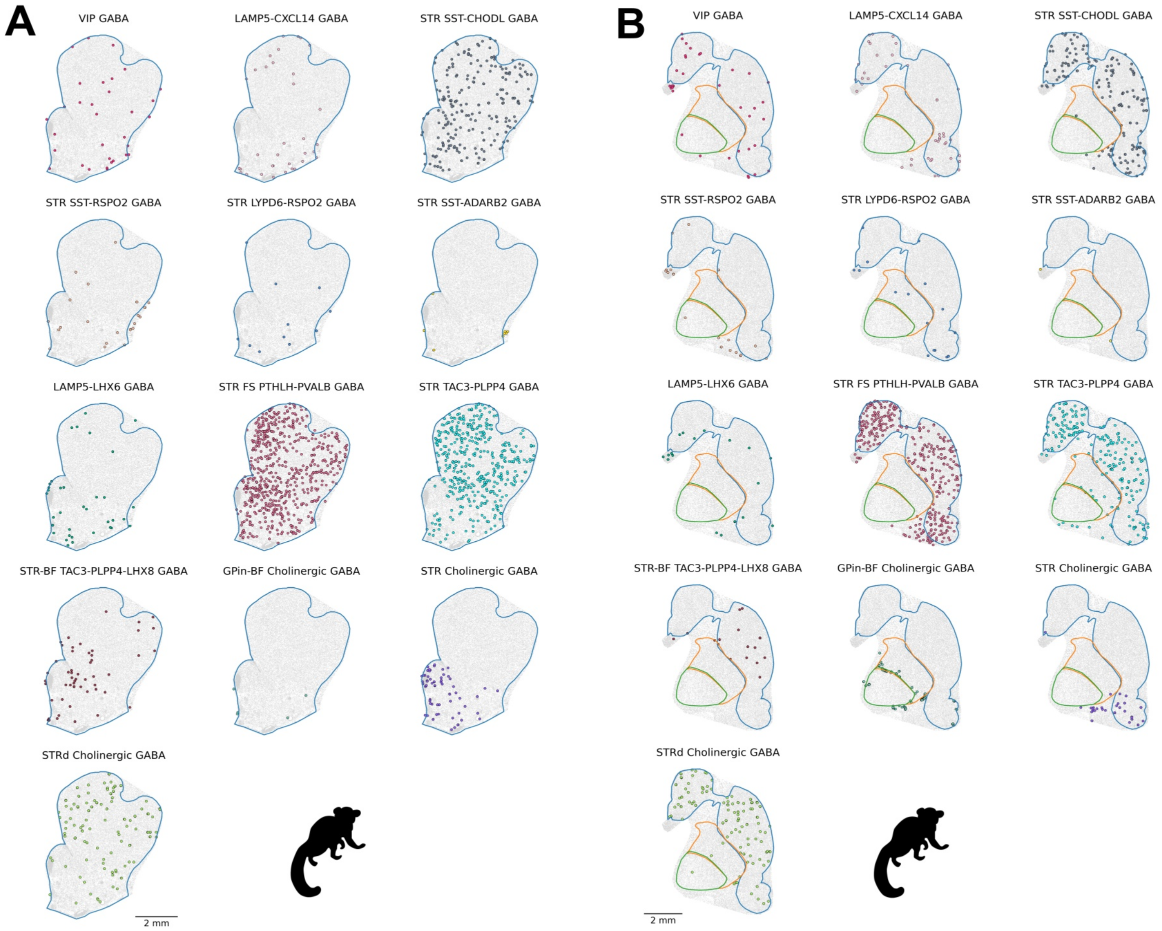


**Supplemental Figure S4:** Spatial distribution of CNU-MGE and CNU-CGE Groups in striatum of marmoset, related to Figure 1

**A.** Representative rostral striatal section. **B.** Representative caudal striatal section. Cells colored by group with all other cells in grey. Regions outlined for reference, STR (blue), GPe (orange), GPi (green).

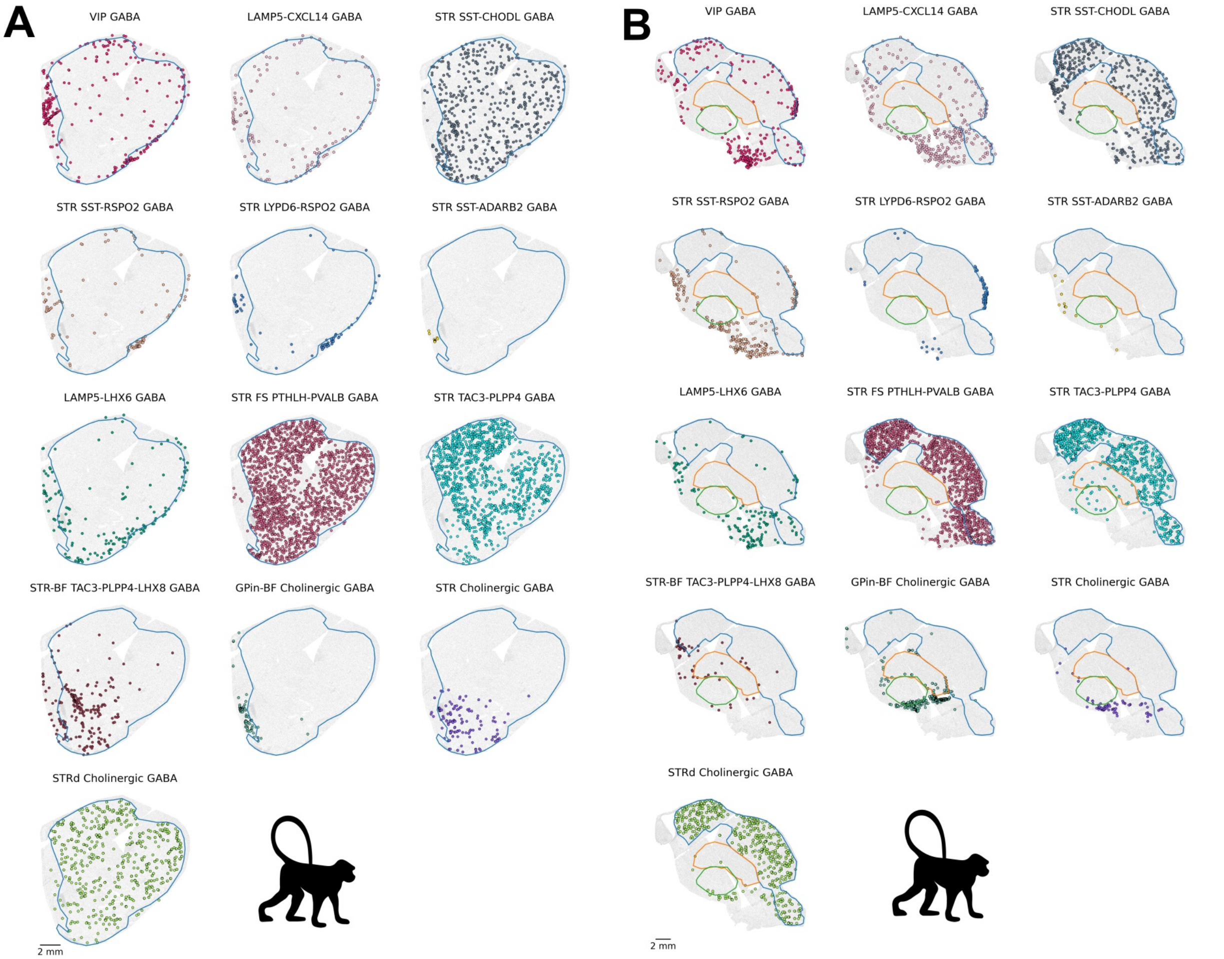


**Supplemental Figure S5:** Spatial distribution of CNU-MGE and CNU-CGE Groups in striatum of macaque, related to Figure 1

**A.** Representative rostral striatal section. **B.** Representative caudal striatal section. Cells colored by group with all other cells in grey. Regions outlined for reference, STR (blue), GPe (orange), GPi (green).

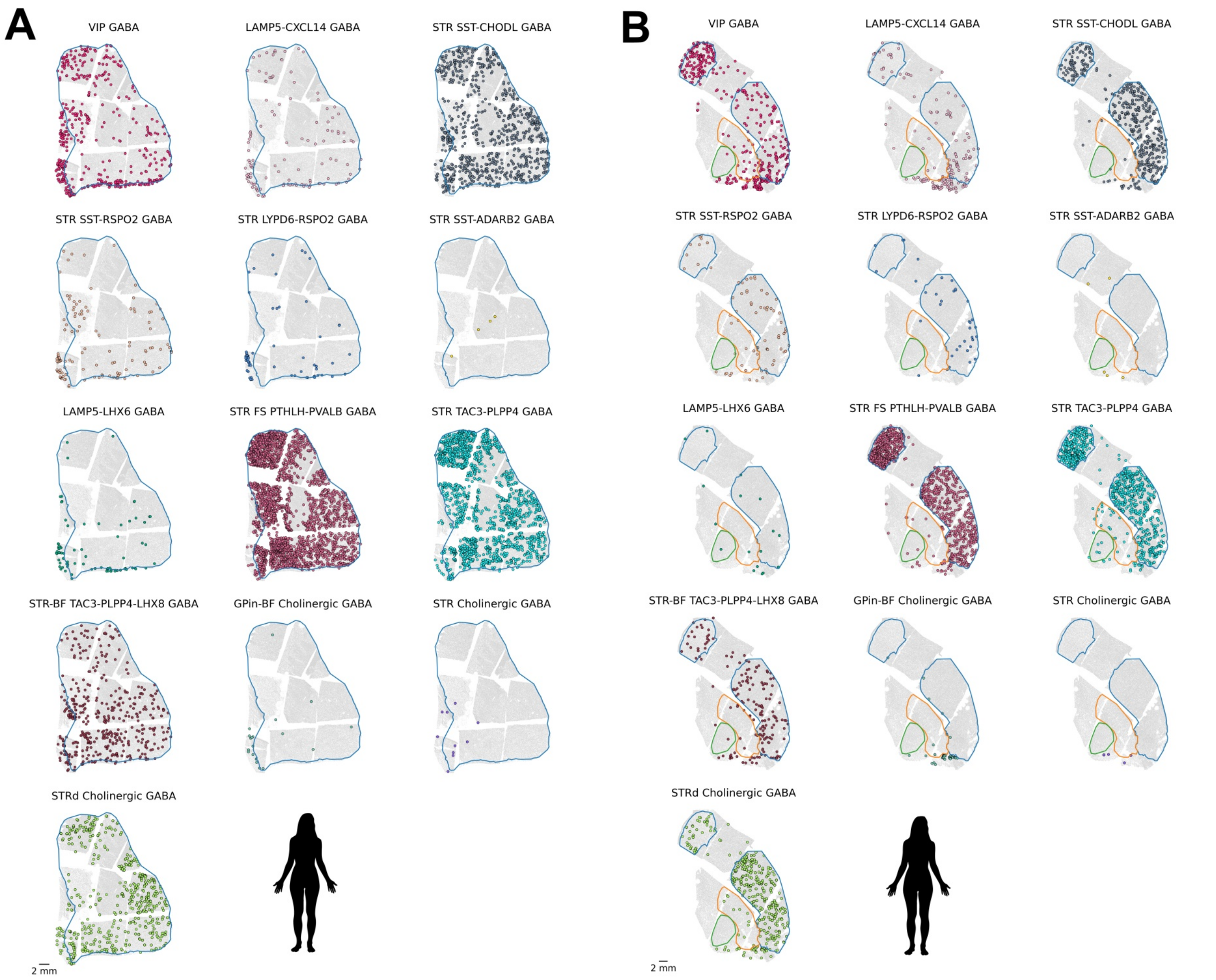


**Supplemental Figure S6:** Spatial distribution of CNU-MGE and CNU-CGE Groups in striatum of human, related to Figure 1

**A.** Representative rostral striatal section. **B.** Representative caudal striatal section. Cells colored by group with all other cells in grey. Regions outlined for reference, STR (blue), GPe (orange), GPi (green).

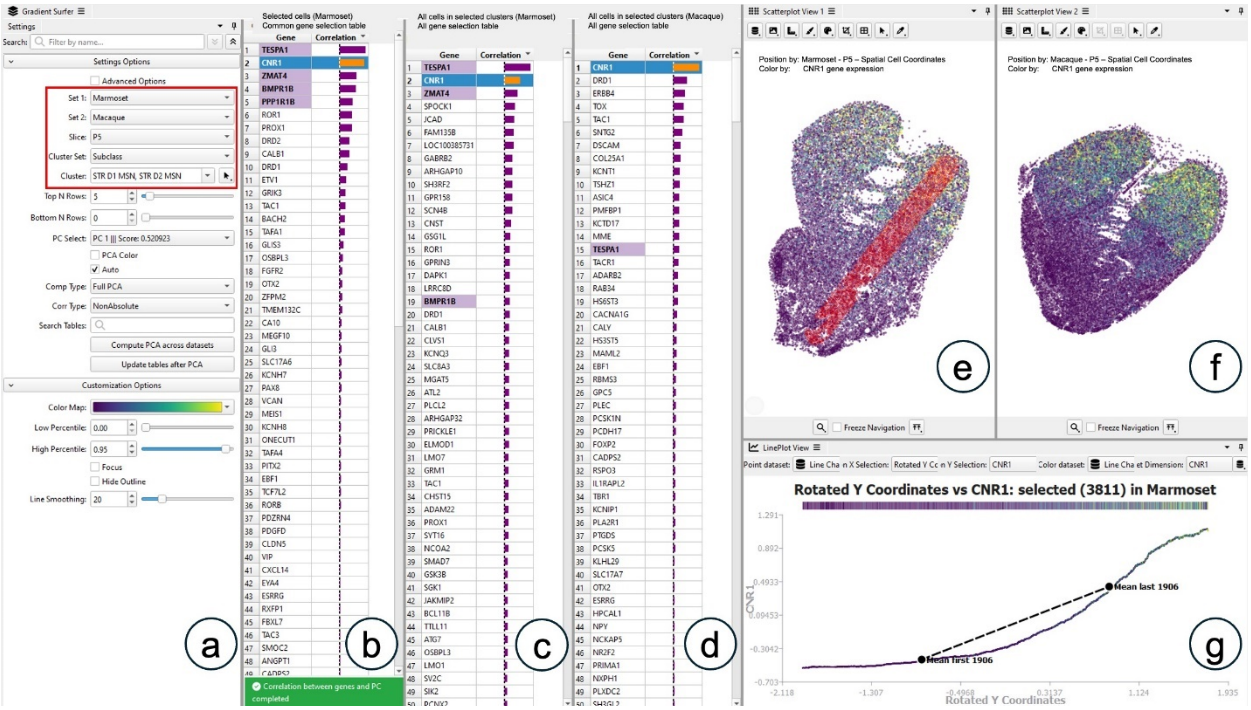


**Supplemental Figure S7:** Cytosplore Gradient Surfer plugin, related to Resource section

Linked-view analysis in Cytosplore Gradient Surfer comparing Marmoset and Macaque datasets, focusing on STR D1 and D2 MSN subclasses. **(a)** Settings panel showing the selected datasets and subclasses. **(b)** Gene correlation table for Marmoset, showing correlations between gene expression and a user-defined spatial gradient of cells along the line in (e). **(c, d)** Gene correlation tables showing correlations between the top principal component and gene expression for all cells in the selected Marmoset (c) and Macaque (d) subclasses. **(e, f)** Spatial maps colored by *CNR1* expression, illustrating the spatial gradients in Marmoset (e) and Macaque (f). **(g)** Line plot showing the *CNR1* expression gradient along the selected spatial axis in Marmoset cells.

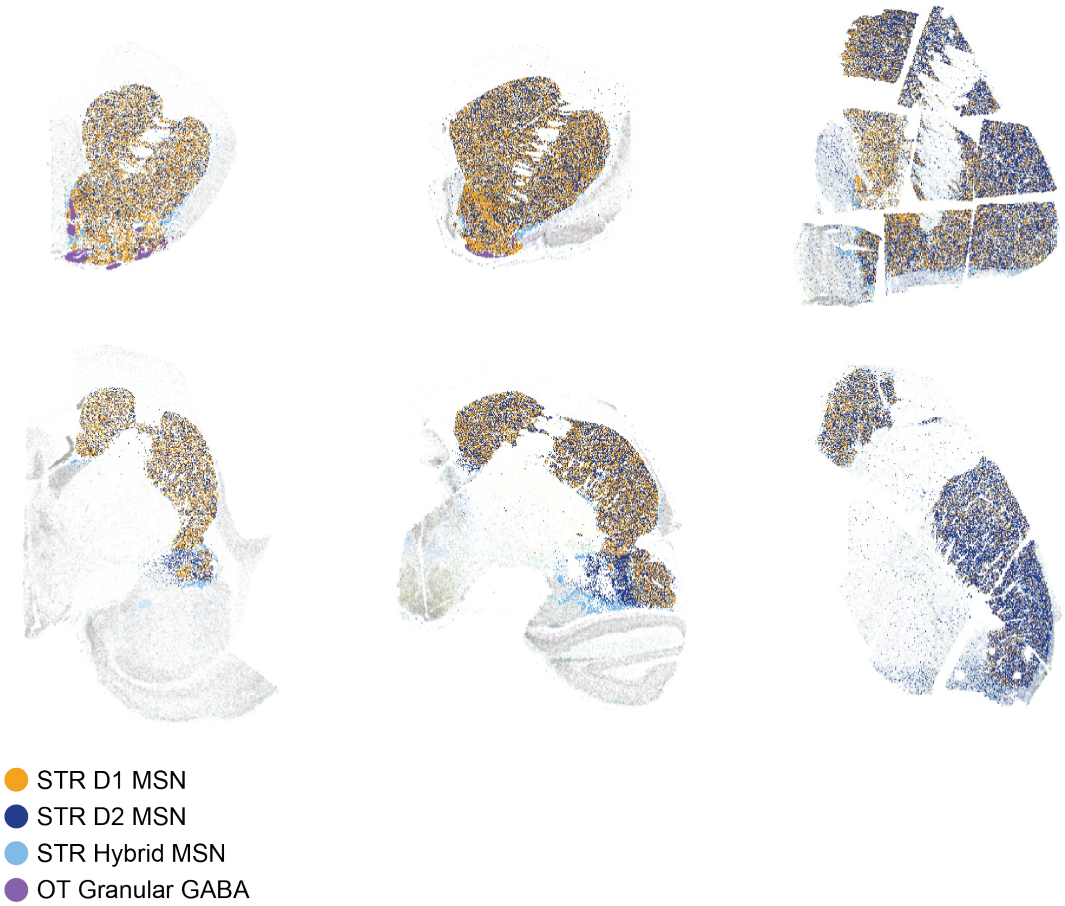


**Supplemental Figure S8:** Neuronal Subclasses in the Striatum, Related to Figure 2. Four dominant subclasses in the striatum, shown at one rostral and one caudal z-plane in each species.

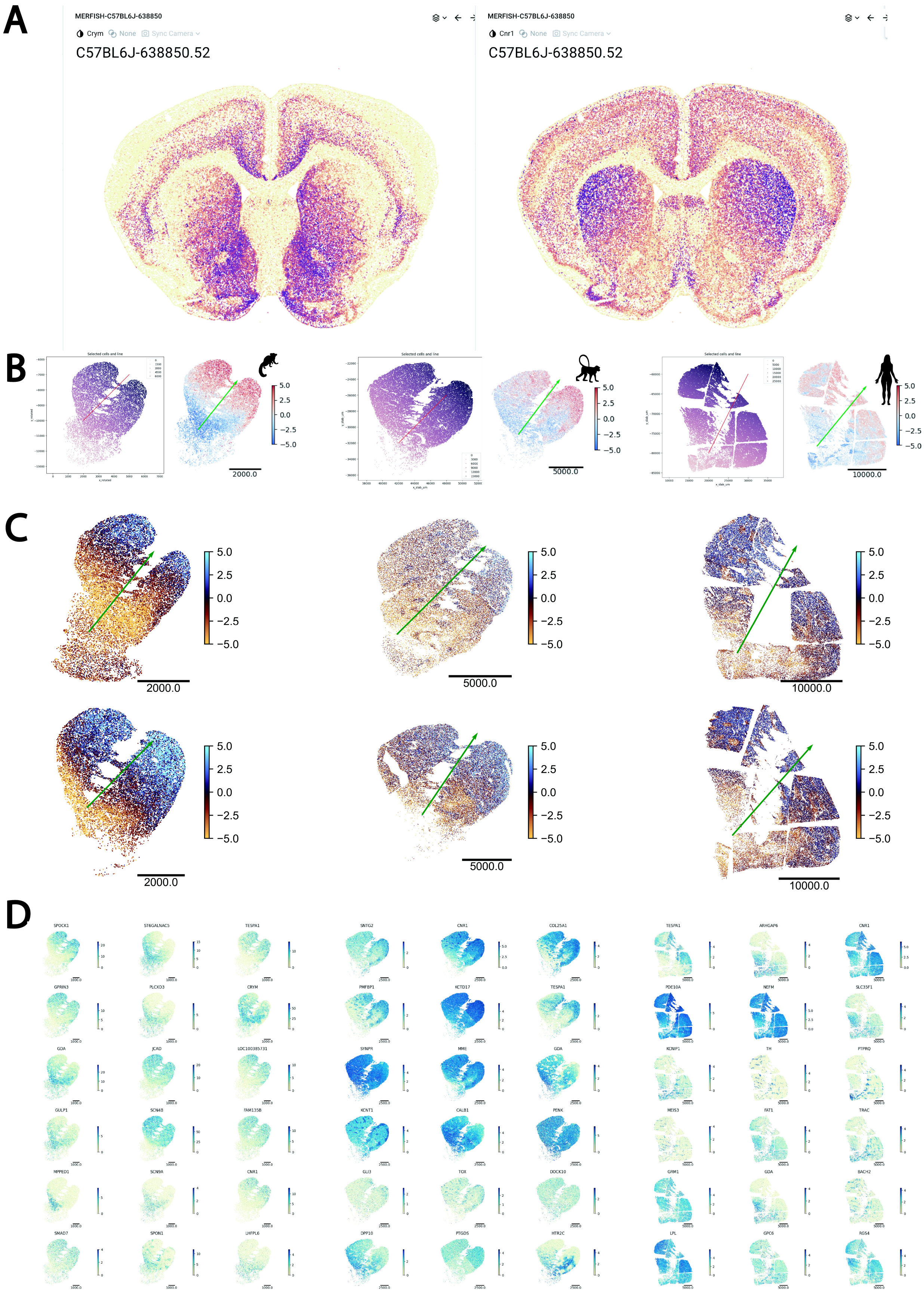
A
MERFISH-C57BL6J-638850
Crym
None
Sync Camera
C57BL6J-638850.52
MERFISH-C57BL6J-638850
Cnr1
None
Sync Camera
C57BL6J-638850.52
B
Selected cells and line
2000.0
5000.0
10000.0
C
2000.0
5000.0
10000.0
D
SPOCK1
ST6GALNAC5
TESPA1
SNTG2
CNR1
COL25A1
TESPA1
ARHGAP6
CNR1
GPRIN3
PLCXD3
CRYM
PMFBP1
KCTD17
TESPA1
PDE10A
NEFM
SLC35F1
GJA
JCAD
LOC100385731
SYNPR
MME
GDA
KCNIP1
TH
PTPRQ
GULP1
SCN4B
FAM135B
KCNT1
CALB1
PENK
MEIS3
FAT1
TRAC
MPPED1
SCN9A
CNR1
GLI3
TOX
DOCK10
GRM1
GDA
BACH2
SMAD7
SPON1
LHFPL6
DPP10
PTGDS
HTR2C
LPL
GPC6
RGS4

**Supplemental Figure S9:** Gene expression gradients in dorsal striatum, related to Figure 3

**A.** Mouse whole brain atlas MERSCOPE data from the ABC Atlas of Crym (left) and Cnr1 (right) detection in striatum, showing the opposing gradients of Crym and Cnr1 described in Stanley et al. **B.** For each species the left panel shows the manually drawn axis of the internal capsule and cells are colored by their projected coordinate along this axis. The right panel shows all STRd MSN cells colored by the principal gradient component. **C.** Additional sampling locations on the rostrocaudal axis for each species. In each species, the top section is more rostral than the section in the main figure, the bottom section is more caudal. **D.** Gradient genes in each species.

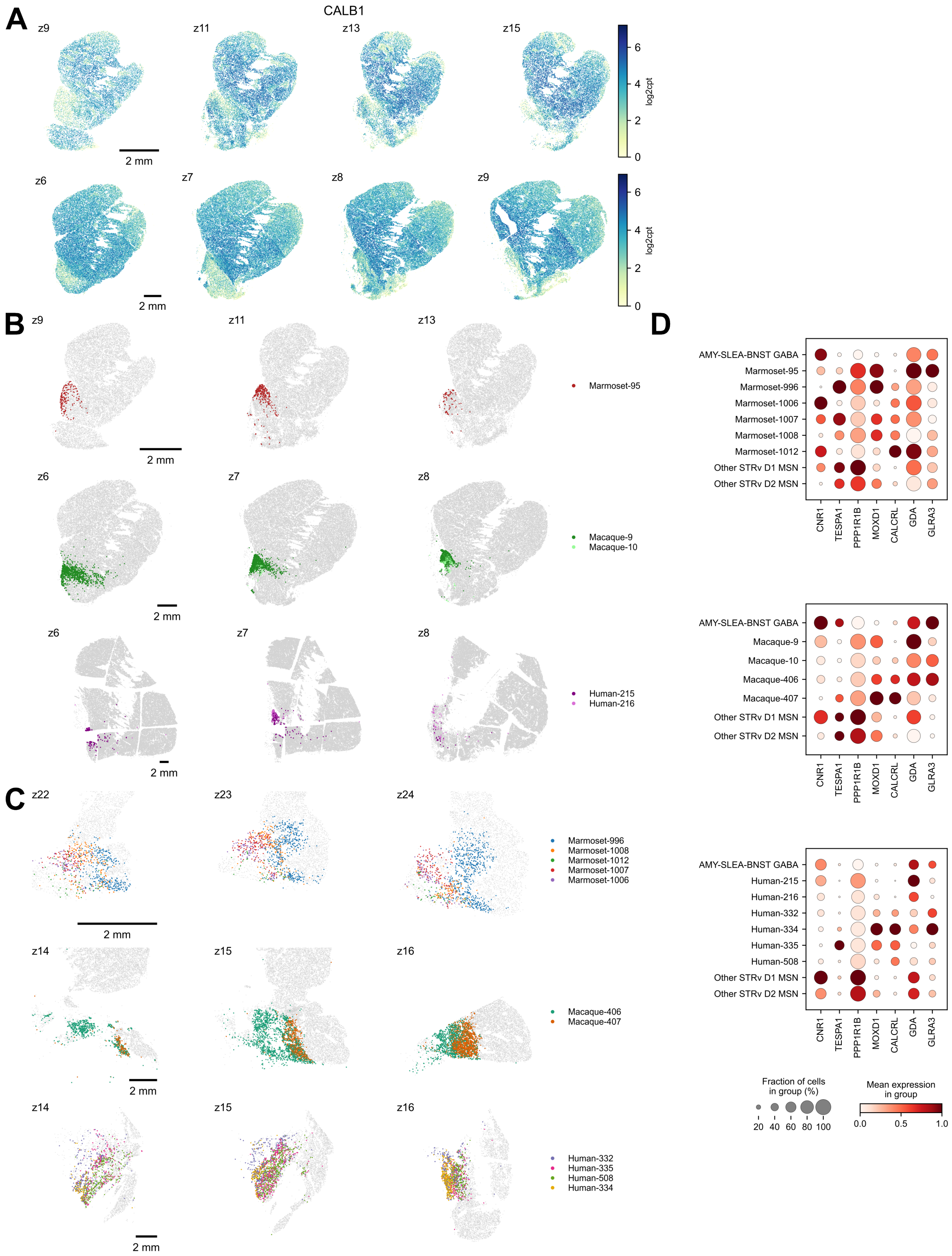


**Supplemental Figure S10:** *CALB1* expression and clusters in ventral striatum, related to Figure 4. **A.** *CALB1* expression for select marmoset (top row) and macaque sections (bottom row). **B.** Selected NACsmd clusters plotted on multiple z planes for marmoset (top row), macaque (middle row), and human (bottom row). MSNs not in the selected clusters are shown in grey. **C.**

Selected AStr clusters plotted on multiple z planes for marmoset (top row), macaque (middle row), and human (bottom row). MSNs not in the selected clusters are shown in grey. **D.** Dot plots of average gene expression in the spatial data for the clusters in B and C.

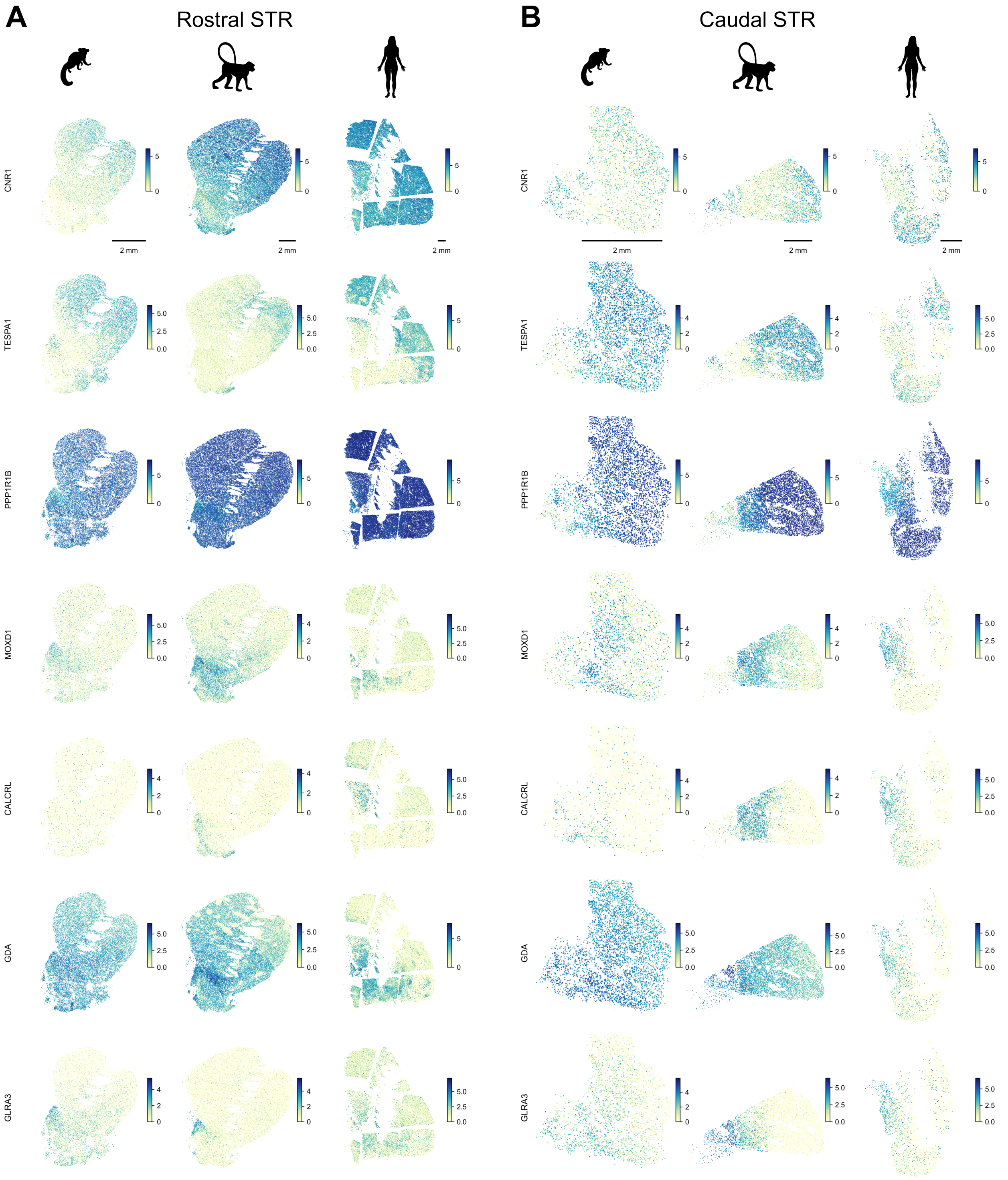


**Supplemental Figure S11:** Spatial gene expression in ventral striatum, related to Figure 4. **A.** Selected genes plotted for a representative rostral STR section for marmoset (left column), macaque (middle column), and human (right column). Log2cpt normalized counts are shown. B**.**

Selected genes plotted for a representative caudal STR section for marmoset (left column), macaque (middle column), and human (right column). Log2cpt normalized counts are shown.

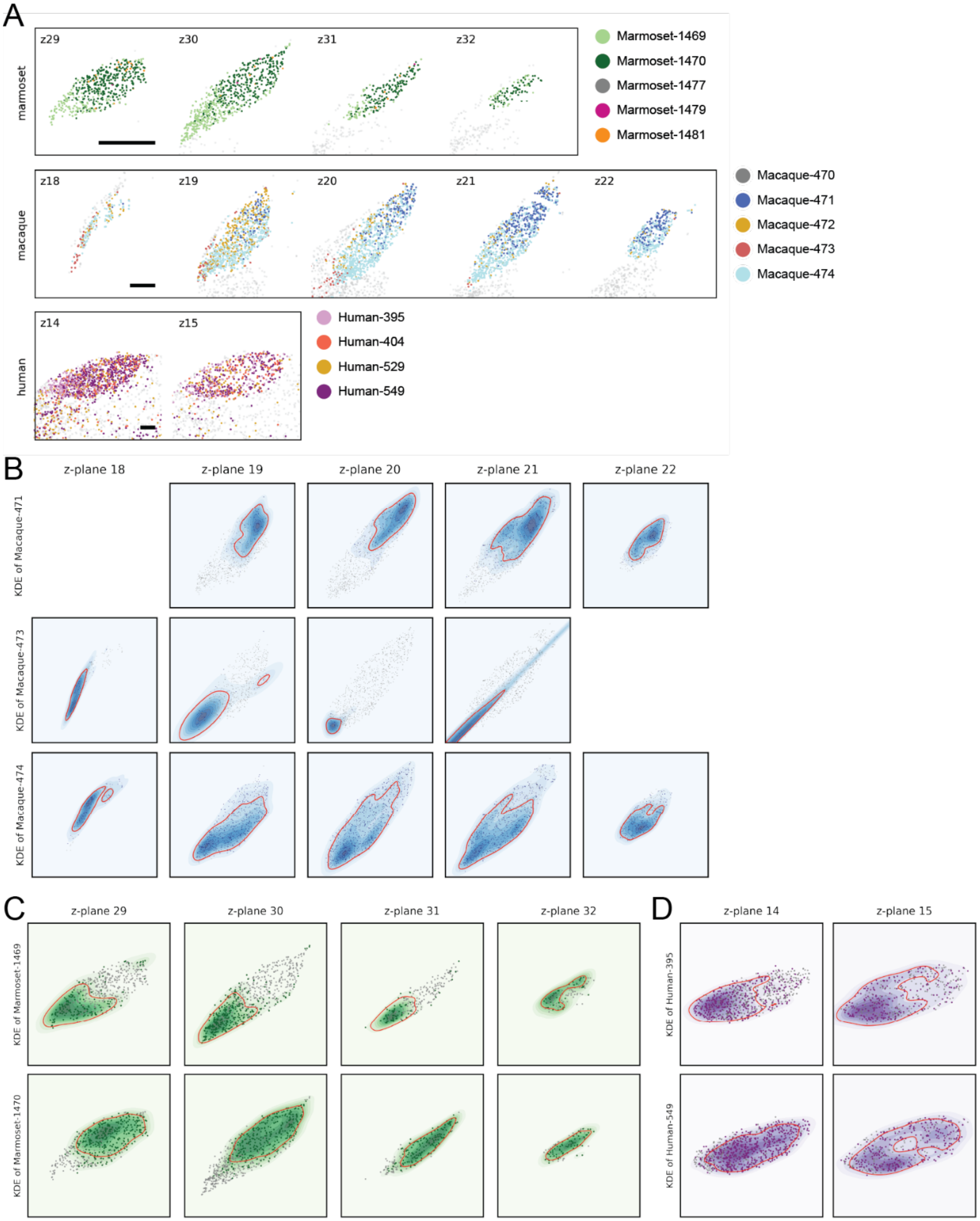


**Supplemental Figure S12:** All subthalamic nucleus (STH) clusters, related to Figure 5.

**A.** All clusters found in the STH in each species. **B-D.** Kernel density estimate (KDE) for selected clusters from **Fig. 5** plotted individually, as contours density maps. The contour that contains 80% of the cells belonging to that cluster is outlined in red. **B.** Macaque. **C.** Marmoset. **D.** Human.

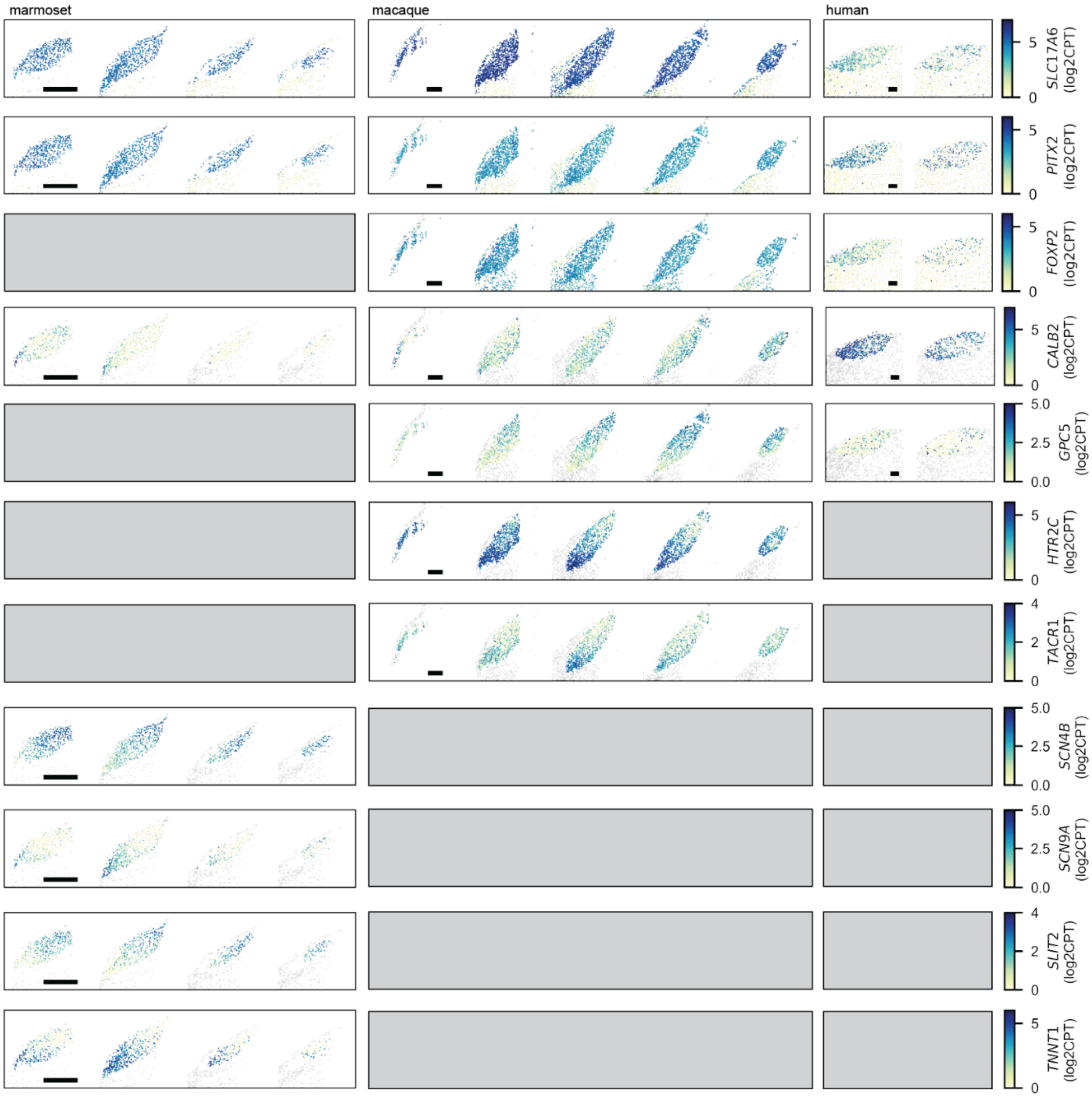


**Supplemental Figure S13:** Spatially variable gene expression in the subthalamic nucleus (STH), related to Figure 5.

Gray boxes indicate the gene was not present in that species' gene panel. All scalebars are 1 mm.

A

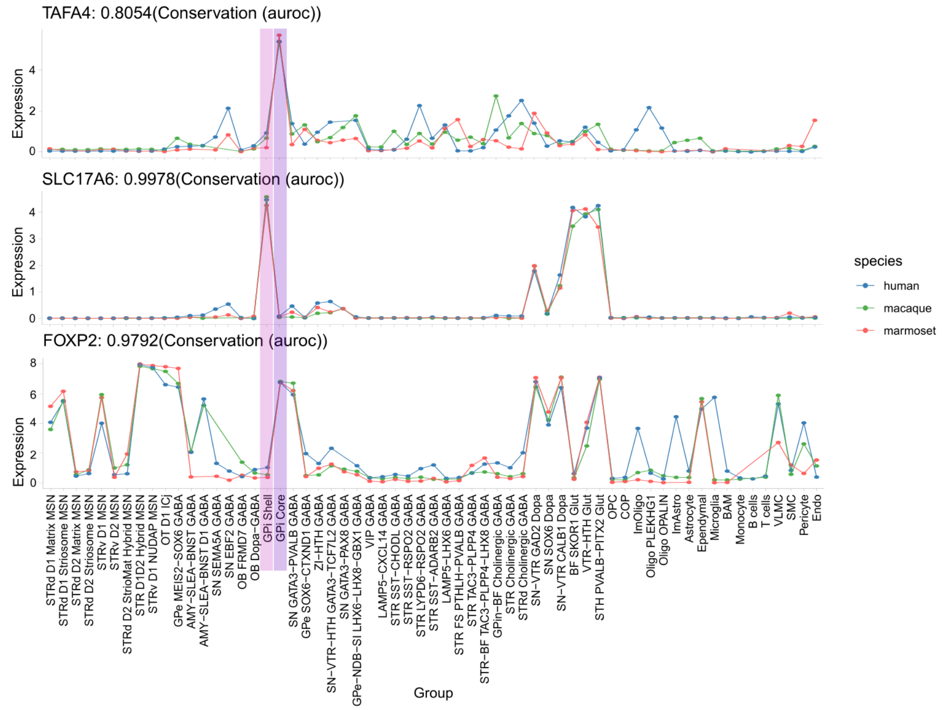


B

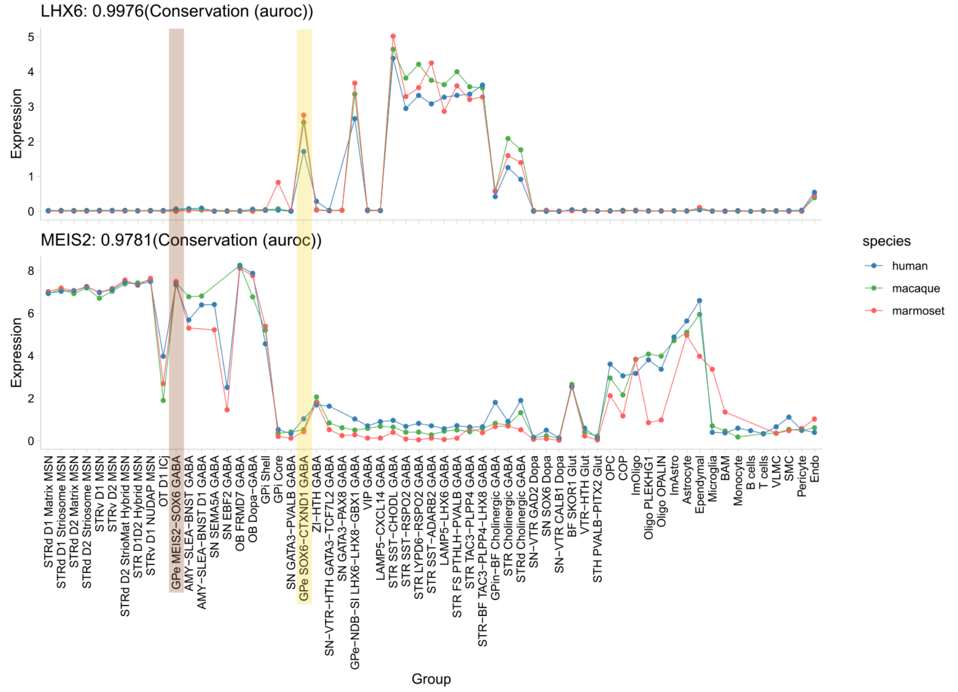


**Supplemental Figure S14:** Expressologs for GP marker genes, related to Figure 6

**A.** *TAFA4* and *FOXP2* are conserved markers for GPi Core (purple) vs i a conserved marker for GPi Shell (pink). **B.** *LHX6* shows conserved, high expression in GPe SOX6-CTXND1 GABA (yellow) vs *MEIS2* which shows conserved, high expression GPe MEIS2-SOX6 GABA (brown).

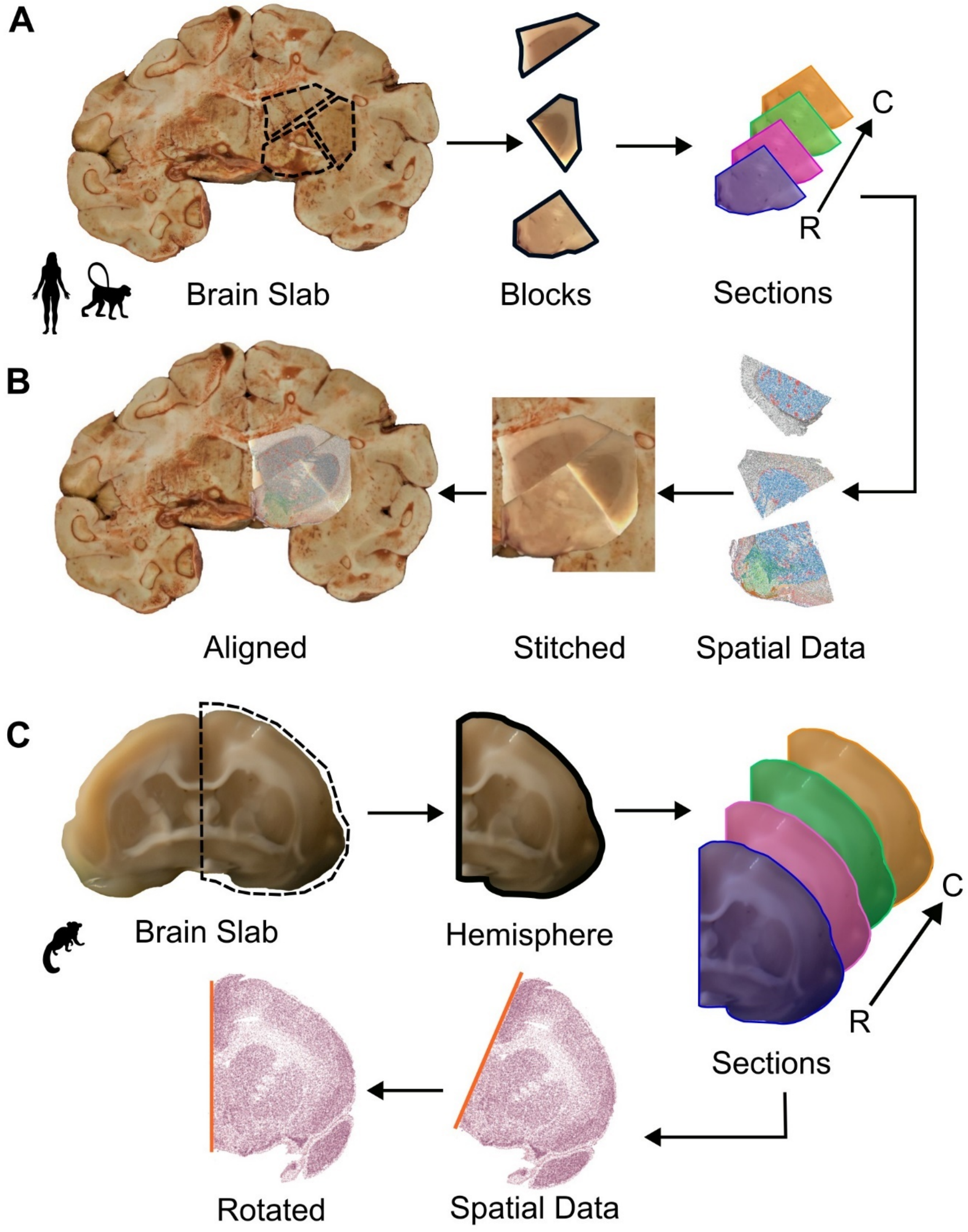


**Supplemental Figure S15:** Tissue processing workflow, related to Methods

**A.** Macaque and human frozen brain slabs were subdivided into blocks approximately 1 $cm^2$ which were sectioned in the rostral to caudal direction for mounting onto MERSCOPE coverslips. **B.** MERSCOPE spatial transcriptomics data from each block was stitched back together guided by block face images of the tissue (middle*)(see Methods*). Stitched spatial data was aligned to slab images (left). **C.** Marmoset brain slabs were hemisected. Slab hemispheres were sectioned in the R-C direction onto Xenium slides. Xenium spatial data was rotated to perpendicular via identification of the midline.
